# *Wolbachia-conferred* antiviral protection is determined by developmental temperature

**DOI:** 10.1101/2020.06.24.169169

**Authors:** Ewa Chrostek, Nelson Martins, Marta S Marialva, Luis Teixeira

**Affiliations:** Institute of Infection, Veterinary and Ecological Sciences, University of Liverpool, United Kingdom; Instituto Gulbenkian de Ciência, Oeiras, Portugal; Institut de Biologie Moléculaire et Cellulaire, Université de Strasbourg, Strasbourg, France; Department for Biomedical Research, University of Bern, Switzerland; Faculdade de Medicina da Universidade de Lisboa, Lisboa, Portugal

**Keywords:** *Wolbachia*, *Drosophila*, virus, temperature, symbiosis, development

## Abstract

*Wolbachia* is a maternally transmitted bacterium widespread in arthropods and filarial nematodes, and confers strong antiviral protection in *Drosophila melanogaster* and other insects. *Wolbachia*-transinfected *Aedes aegypti* are currently being deployed to fight transmission of dengue and Zika viruses. However, the mechanism of antiviral protection and factors influencing it are still not fully understood. Here we show that temperature modulates *Wolbachia*-conferred protection in *Drosophila melanogaster*. Temperature after infection directly impacts *Drosophila* C virus replication and modulates *Wolbachia* protection. At higher temperatures virus proliferates more and is more lethal, while *Wolbachia* confers lower protection. Strikingly, host developmental temperature is a determinant of *Wolbachia*-conferred antiviral protection. While there is a strong protection when flies are raised from egg to adult at 25°C, the protection is highly reduced or completely abolished when flies develop at 18°C. However, *Wolbachia*-induced changes during development are not sufficient to limit virus-induced mortality, as *Wolbachia* is still required to be present in adults at the time of infection. This developmental effect is general, since it was present in different host genotypes, *Wolbachia* variants and upon infection with different viruses. Overall, we show that *Wolbachia*-conferred antiviral protection is temperature dependent, being present or absent depending on the environmental conditions. This interaction likely impacts *Wolbachia*-host interactions in nature and, as a result, frequencies of host and symbionts in different climates. Dependence of *Wolbachia*-mediated pathogen blocking on developmental temperature could be used to dissect the mechanistic bases of protection and should be considered by programmes deploying *Wolbachia* as an antiviral agent in the field.

**Significance Statement:** Insects are often infected with beneficial intracellular bacteria. The bacterium *Wolbachia* can protect insects from pathogenic viruses. This effect can be used to prevent transmission of dengue and Zika viruses by *Wolbachia*-infected mosquitoes. To deploy *Wolbachia* in the field successfully and understand the biology of insects in the wild we need to discover which factors affect *Wolbachia*-conferred antiviral protection. Here we show that the temperature in which insects develop from eggs to adults can determine presence or absence of antiviral protection. The environment, therefore, influences this insect-bacterium interaction. Our work may help to provide insights into the mechanism of viral blocking by *Wolbachia* and inform programs using *Wolbachia* in mosquito-borne disease control.

## Main Text

### Introduction

*Wolbachia* is a maternally inherited intracellular bacterium infecting a wide range of arthropods and some nematodes. This endosymbiont induces strong phenotypes in many of its hosts, which may contribute to its success in invading and being maintained in host populations. *w*Mel, the *Wolbachia* strain present in *Drosophila melanogaster*, confers a strong protection to a wide range of RNA viruses, which can be a fitness benefit in nature (1, 2). This protection extends to non-native *Wolbachia*-hosts associations, including mosquito vectors of human disease (3). Recently, *Wolbachia* has become one of the most promising approaches to control dengue and Zika viruses in the wild. It has been shown that the releases of *Wolbachia*-infected *Aedes aegypti* can reduce the number of dengue cases in both, endemic and non-endemic areas (4–7). Although molecular mechanisms of *Wolbachia*-conferred antiviral protection are not yet known, identification of factors influencing protection can contribute to understanding the association of *Wolbachia* and hosts in natural population, as well as guide *Wolbachia*-based field interventions.

*Wolbachia-*conferred antiviral protection is influenced by host and bacterial genetics. Different *Wolbachia* strains in the same host genetic background vary in protection (8–12). In general, differences in *Wolbachia* titres are correlated with differences in protection, with higher titres conferring higher antiviral protection (8–12). However, some *Wolbachia* strains do not provide protection despite high titres (13). Host genetic variation can also contribute to the strength of *Wolbachia*-conferred protection, as seen in *Aedes aegypti* (14).

Environmental factors also affect this symbiont-mediated protection, either through modulation of *Wolbachia* titre or by titre-independent mechanisms. Host diet rich in cholesterol reduces antiviral protection in *Drosophila melanogaster* (15). Similarly, antibiotic treatment reducing *Wolbachia* titre reduces the antiviral protection (16). Finally, temperature was shown to modulate *Wolbachia-* conferred protection to parasites in an engineered *Wolbachia*-host association; in *Anopheles stephensi*, temperature and somatic *Wolbachia* infection determined *Plasmodium* titre in mosquitoes (17). On the other hand, in *Aedes aegypti* stably transinfected with *w*Me*l*, the comparison of a constant and two fluctuating temperature regimes concluded, that the antiviral protection is robust across conditions and unlikely to be compromised in the antiviral field trials (18). However, different temperatures regimes affect dengue transmission independently of *Wolbachia* (19).

Despite two studies tackling temperature effect on *Wolbachia*-conferred protection in artificial associations, the impact of temperature on antiviral protection in natural *Wolbachia-*host symbiosis remains unknown. Yet, it is known that temperature can affect the interaction between *Wolbachia* and *D. melanogaster*. For instance, higher temperature leads to higher proliferation and cost of the pathogenic variant *w*MelPop (20–22). Also, flies with different *w*Mel variants prefer different temperatures, probably related with cost of these variants to the host (23). Geographic distribution of *Wolbachia* in *D. melanogaster* populations could be explained by their relative fitness effects at varying thermal conditions (24, 25). Antiviral protection could be a temperature-dependent fitness benefit, which would be balanced against the potential cost of highly protective, high titre symbionts.

Here we tested how temperature affects *Wolbachia* protection to viruses in natural *Wolbachia-Drosophila* associations. First, we asked how different virus doses and post-infection temperatures affect *Wolbachia* densities and antiviral protection. Then, we dissected the effect of developmental temperature on *Wolbachia*-carrying *Drosophila melanogaster*. We show strong interaction between this environmental variable and *Wolbachia*-conferred protection against viruses.

### Results

To test how different infection temperatures affect *Wolbachia*-conferred protection to *Drosophila* C virus (DCV), we used the *Drosophila melanogaster* Drosdel *w*^*1118*^ isogenic line (*iso)* carrying a natural *Wolbachia* variant, *w*MelCS_b (*Wolb*+), and a matching *Wolbachia-*free control (*Wolb*-) (1, 8). Flies were raised at 25°C and 3-6-day-old flies were challenged with serial dilutions of DCV and subsequently placed at either 18°C or at 25°C (Fig. 1A, B). Virus-induced mortality was higher at 25°C than 18°C at all doses except the lowest one, where there is almost no mortality associated with the infection (Temperature effect, *p* ≤ 0.001 for all comparisons, except with infection dose of 10^5^ TCID_50_/ml). Importantly, *Wolbachia* protection against DCV varies with temperature (*Wolbachia*\*Temperature interaction effect, *p* < 0.001), and it is stronger when temperature of infection is 18°C (Cox hazard ratio (CHR) = -1.84, *p* < 0.001) than 25°C (CHR = -0.75 *p* < 0.001, at 18°C). Overall, the temperature of infection affects the survival of the flies and *Wolbachia* protects more at lower infection temperature (Fig. 1A,B).

**Figure 1.**
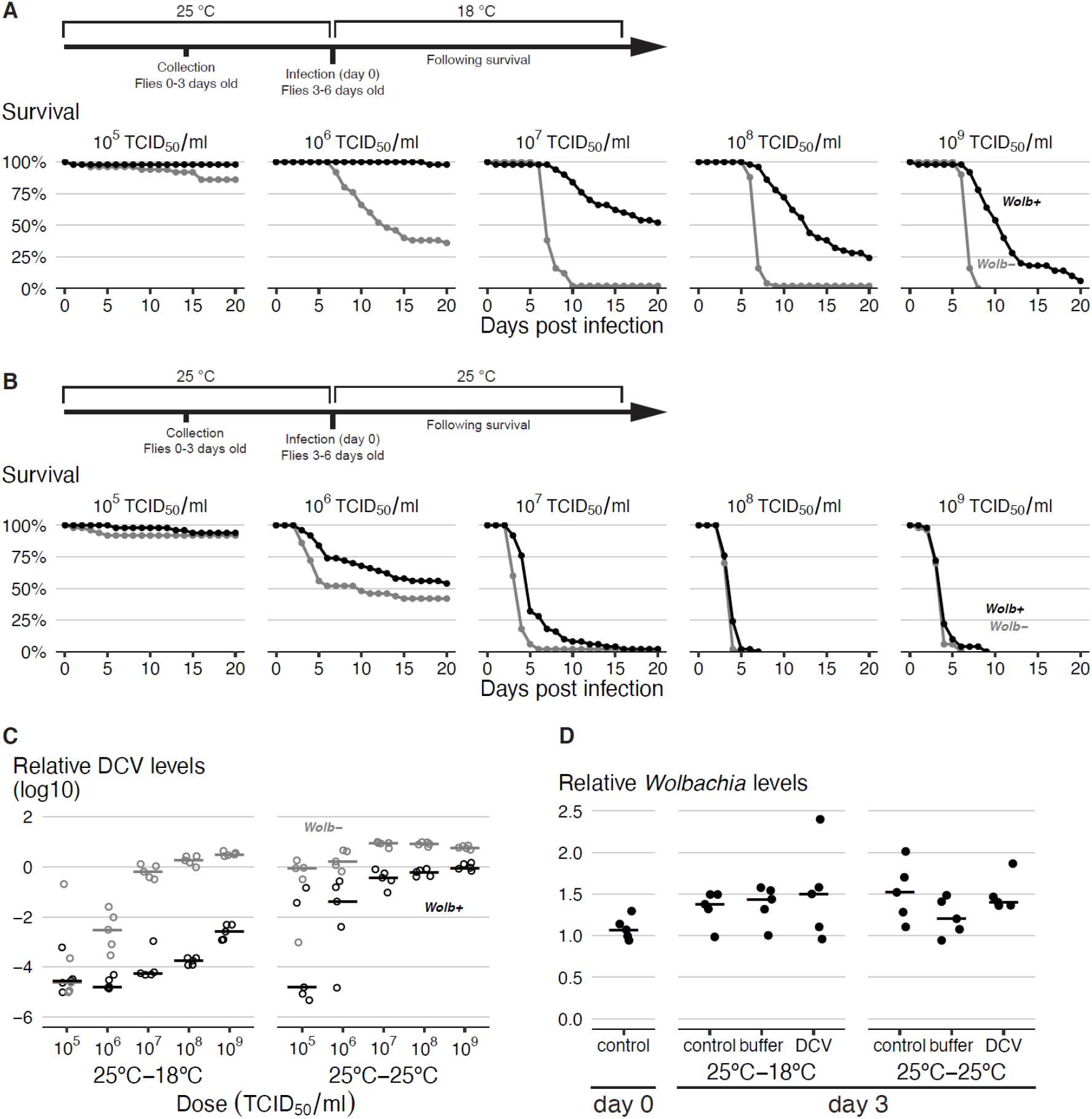
Post-infection temperature modulates the strength of *Wolbachia*-conferred antiviral protection. (A,B) Survival curves of flies infected at different temperatures, with the schemes of the experimental designs shown above. *Wolb+* and *Wolb-* flies, 50 per *Wolbachia* status and condition, were raised at 25°C, infected with different doses of DCV and placed at either 18°C (A) or 25°C (B) after DCV infection. Mortality was recorded daily. C) DCV titres in DCV-infected flies 3 dpi measured by RT-qPCR. Flies were kept at 25°C before infection and either at 18°C or 25°C after infection. (D) *Wolbachia* levels measured by qPCR at the day of infection (3-6 days old), or 3 days after in unchallenged (control), buffer- or DCV-challenged (10^7^ TCID_50_/ml) flies. Flies were kept at 25°C before and placed at either 18°C or 25°C after DCV infection. In (C) and (D) each dot is a sample of ten flies and horizontal lines are medians.

To test if the interaction of *Wolbachia* protection with temperature is also reflected in viral loads, we measured viral titres in flies three days post infection (dpi) by quantitative reverse transcription PCR (RT-qPCR) (Fig. 1C). We detected a strong interaction between *Wolbachi*a presence, temperature of infection, and dose of the virus (*p* < 0.001). At the lower temperature *Wolbachia* confers more resistance at higher viral doses, as viral titres stay very low at lower doses in both *Wolb+* and *Wolb-* flies. At the higher temperature, it confers more resistance at the lower doses, as virus titres are very high in *Wolb+* and *Wolb-* flies exposed to high virus doses. The mean viral load was higher at 25°C than at 18°C (*p* < 0.001) and was lower in the presence of *Wolbachia* (*p* < 0.001). On average, *Wolbachia* induces higher resistance at 18°C than at 25°C. There was a 550-fold reduction in viral load at 18°C (*p* < 0.001) and 50-fold reduction at 25°C (*p* < 0.001), producing approximately 11-fold difference between these two conditions (*p* = 0.003). These results show that the strength of *Wolbachia*-conferred protection to DCV, in terms of both, survival and viral loads, depends on the temperature of infection. Protection is higher at a lower temperature. Since post-infection temperature affects viral infection independently of *Wolbachia*, the lower capacity of *Wolbachia* to protect at higher post-infection temperatures may be related with a higher replication of the virus.

Antiviral protection is also usually positively correlated with *Wolbachia* levels (8, 9, 11, 16, 26, 27), however, so we tested *Wolbachia* levels at the day of infection and after three days in DCV-infected, buffer-pricked (control), or unmanipulated flies, at both temperatures (Fig. 1D). *Wolbachia* levels were not affected by virus, buffer or temperature of infection (*p* > 0.425 for effect of temperature or treatment). Thus, the difference in protection is likely independent of *Wolbachia* levels, as these are not significantly affected within three days of the viral challenge.

We extended our analysis by testing how pre-infection temperature affects *Wolbachia* protection (Fig. 2). We followed survival of infected flies in all four possible combinations of pre- and post-infection temperatures of 18°C and 25°C (Fig. 2A, B). There is a strong interaction between pre-infection temperature and *Wolbachia* (*p* = 0.009). Remarkably, in flies raised at 18°C *Wolbachia* does not protect against DCV infection (*p* = 0.207), while in flies raised at 25°C it does (CHR = -1.38, *p* < 0.001), irrespectively of the post-infection temperature. Also, in the absence of *Wolbachia* the pre-infection temperature has no effect on the survival after DCV infection (*p* = 0.276). This strong interaction between *Wolbachia*-conferred protection to DCV and pre-infection temperature is also reflected in DCV loads (Fig. 2C, S1, *p* < 0.001). On average, *Wolbachia* reduced viral titres 14-fold at pre-infection temperature of 18°C and 1900-fold at 25°C (difference *p* < 0.001). In these assays, pre-infection temperature also did not affect viral loads in the absence of *Wolbachia* (*p* = 0.534). Therefore, in contrast to post-infection temperature, pre-infection temperature does not directly impact virus performance and modulates only *Wolbachia* protection. In summary, in flies raised at lower temperature *Wolbachia* confers a reduced resistance to DCV in terms of viral titres and no protection in terms of survival.

**Figure 2.**
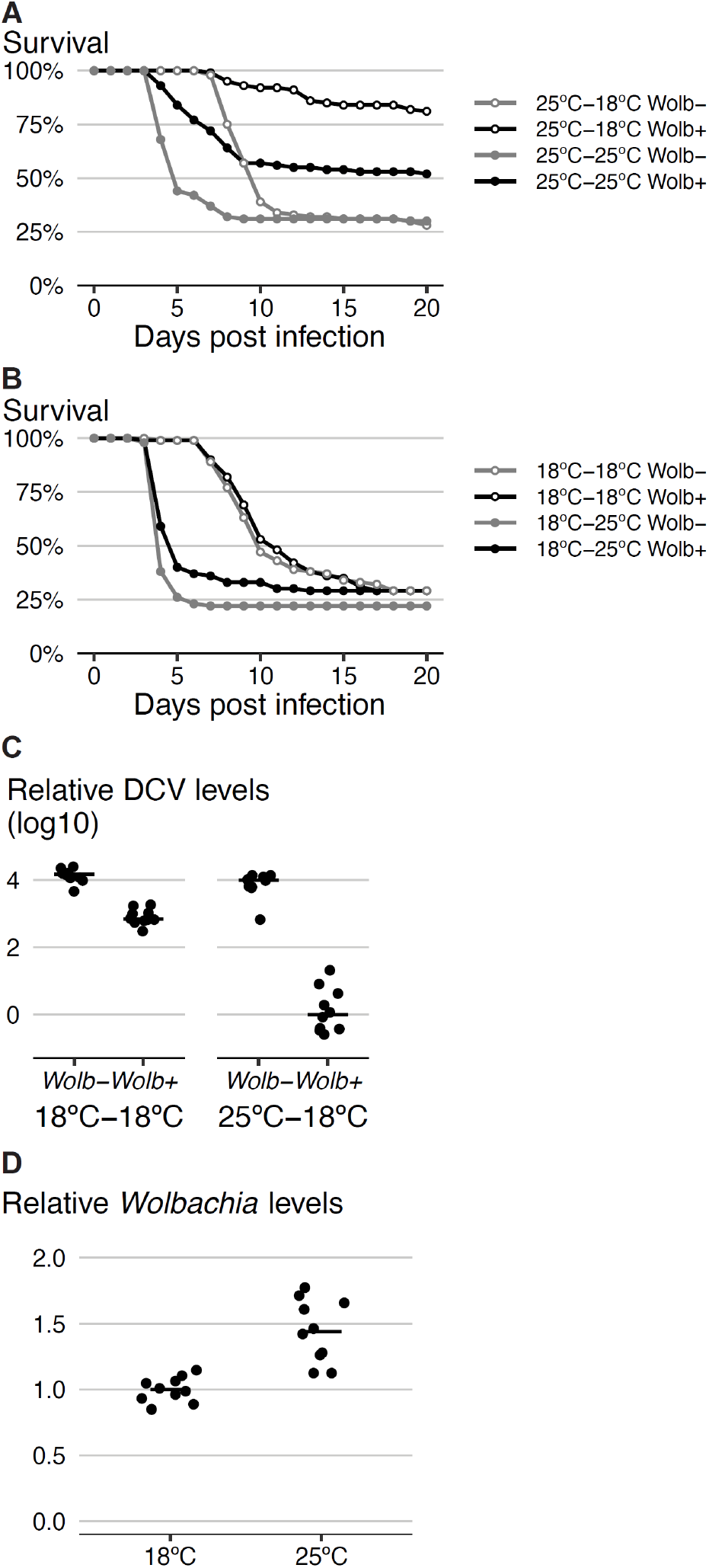
Pre-infection temperature determines *Wolbachia*-conferred antiviral protection. (A,B) *Wolb+* and *Wolb-* flies, fifty per *Wolbachia* status per condition, were infected with DCV (10^8^ TCID_50_/ml) and checked for survival every day. Flies were raised and kept at 25°C (A) or 18°C (B) before DCV infection and placed at either 18°C or 25°C after infection. (C) DCV titres in flies 3 dpi measured by RT-qPCR. Flies were raised and kept at either 18°C or 25°C before infection and at 18°C after infection. Replicate of this experiment is shown in Fig. S1. (D) *Wolbachia* levels measured by qPCR in 3-6 days old flies raised at either 18°C or 25°C. For (C) and (D) each dot is a sample of ten flies. Horizontal lines are medians of the samples.

Next, we measured *Wolbachia* levels in flies raised at 18°C and 25°C (Fig 2D). Flies raised at 18°C had approximately 33% less *Wolbachia* than the flies raised at 25°C (*p* < 0.001). Although this difference may contribute to the difference in protection, a larger, 50% difference in *Wolbachia* titres between *w*Mel- and *w*MelCS-harbouring flies did not abolish the protection (8). Thus, differences in *Wolbachia* titres do not seem to fully explain the difference in protection between flies raised at different temperatures.

To test the robustness of pre-infection temperature effect on *Wolbachia-*conferred protection, we tested flies with distinct host genetic backgrounds, Aljezur1 and Oregon-R W-20 (1), each harbouring its original *w*Mel-like *Wolbachia* strain (Fig. 3A,B and S2A,B). In agreement with the previous results, there was a significant effect of pre-infection temperature on *Wolbachia*-conferred antiviral protection against DCV in both lines (*p*<0.001 for both). *Wolbachia* in Aljezur1 line conferred protection only when the pre-infection temperature was 25°C (*p* = 0.763 for 18°C, and CHR = -1.19, *p* < 0.001 for 25°C). In the Oregon-R W-20 line, *Wolbachia* still conferred some protection when pre-infection temperature was 18°C, although it was significantly less than at 25°C (CHR = -1.21, *p* < 0.001 for 18°C, and CHR = -3.02, *p* < 0.001 for 25°C, CHR difference = 1.8, *p* < 0.001). Next, we investigated if the pre-infection temperature effect was specific to DCV or if it was also present upon challenge with another RNA virus, Flock house virus (FHV) (Fig. 3C and S2C), There was an interaction between *Wolbachia*, pre-infection temperature, and dose in this assay (*p* = 0.013). *Wolbachia* protection was significantly higher at flies raised at 25°C for all doses of virus except for the lowest one (contrasts of CHR < -2.84, *p* < 0.001 for 10^7^ – 10^9^ TCID_50_/ml). At the lowest dose, 10^6^ TCID_50_/ml, there was an overall low mortality of flies and the difference in survival was not statistically significant (contrast of CHR = -1.00 but *p* = 0.421). The pre-infection temperature had no effect on survival to FHV in the absence of *Wolbachia* (*p* = 1, in all comparisons), as observed for DCV. In conclusion, the strong influence of pre-infection temperature on *Wolbachia*-conferred protection is a general effect observed in different host genetic backgrounds, *Wolbachia* variants and with the different viruses used.

**Figure 3.**
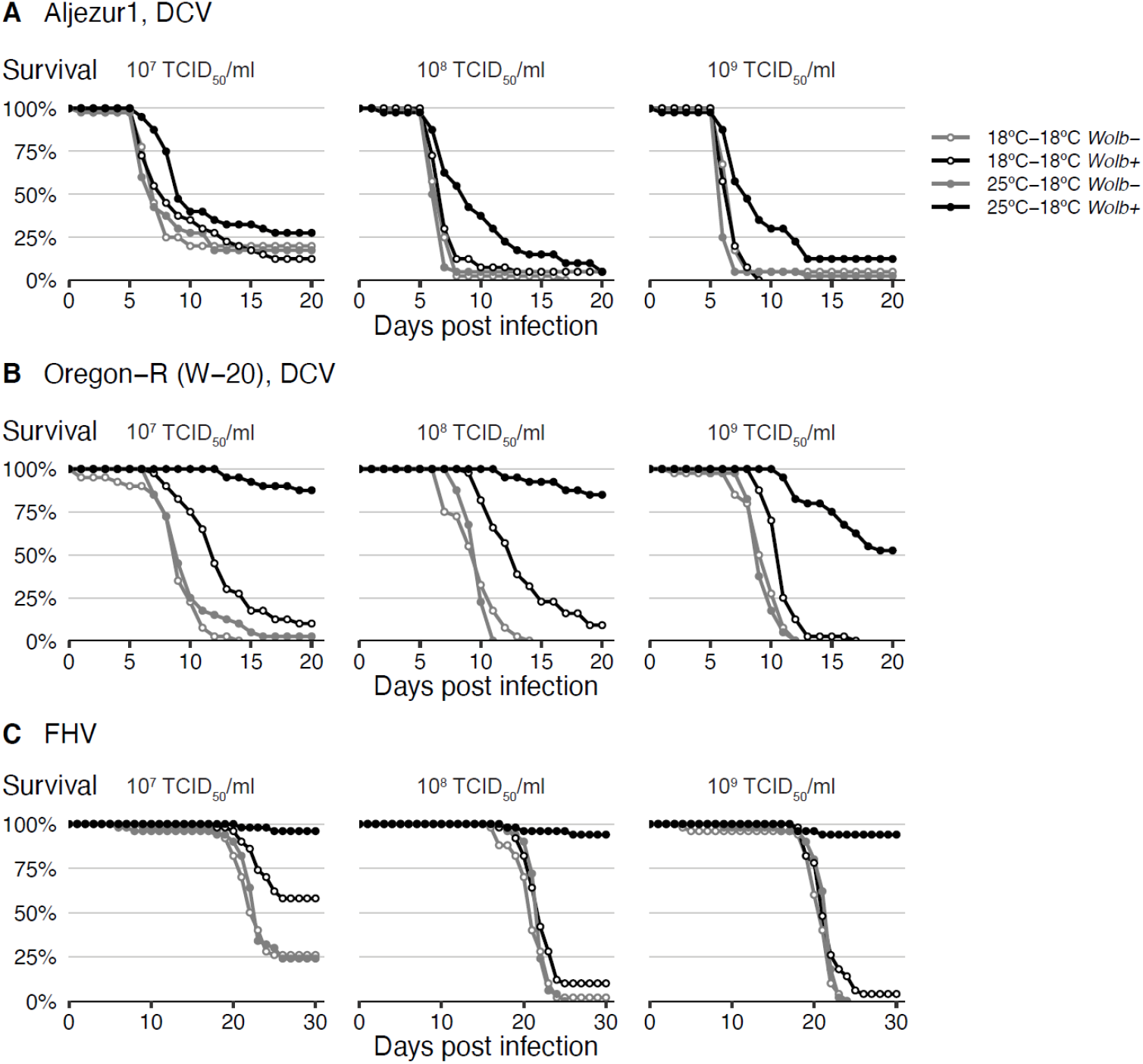
Pre-infection temperature determines *Wolbachia*-conferred antiviral protection in different *Wolbachia* and *Drosophila* genotypes and against different viruses. (A) Aljezur 1 and (B) Oregon-R (W-20) *Wolb+* and *Wolb-* flies, fifty per *Wolbachia* status per condition, were pricked with three dilutions of DCV. (C) *w*^1118^ *iso* flies, fifty per *Wolbachia* status per condition, were infected with three doses of FHV. Flies were kept at 25°C or 18°C before the infection, and at 18°C after the infection. Mortality was recorded daily. A replicate of this experiment is shown in Fig. S2.

Since there is a strong interaction between protection and constant pre-infection temperature, we asked if protection exists when temperature is cycling daily between 18°C and 25°C (Fig. S3). *Wolbachia* protects against DCV infection at the cycling temperature (CHR = -0.67, *p* = 0.014). This protection seems to be intermediate between the protection seen in flies developed at 18°C or 25°C. This result shows that in daily cycling temperatures, which occur in natural conditions, *Wolbachia* can protect against viruses.

In the above protocols, flies developed from egg to adult at a given temperature and after collection they were aged for three more days at the same temperature, before infection with viruses. To dissect at which of these stages, development or aging, temperature influences *Wolbachia* protection, we tested the four possible combinations of the two different temperatures for these two stages (Fig. S4). Developmental temperature strongly influenced *Wolbachia* protection, which was higher at 25°C (CHR = -1.16, *p* < 0.001). Aging the flies at 25°C also interacted with *Wolbachia* protection, but increased it only slightly compared to aging at 18°C (CHR = -0.431, *p* = 0.011). Therefore, *Wolbachia* protection to viruses is mainly dependent on the temperature of development form egg to adult.

Since pre-infection events are crucial for *Wolbachia*-conferred resistance we asked how fast the difference in virus titres arise between flies with and without *Wolbachia* (Fig. 4A). We observed a *Wolbachia*-induced resistance of approximately 120-fold as soon as we can detect viral RNA, at 12h after infection, (*p* < 0.001 at 12h, and *p* < 0.002 for all posterior time points in Fig 4A and S5A). Thus, *Wolbachia* lowers viral titres very early in the course of DCV infection. We observed a similar pattern of early FHV blocking by *Wolbachia*, from day one post infection onwards, but with smaller viral titre differences (*Wolbachia*-infected flies had 2.3 to 5.5 times less FVH from the first day onwards, *p* > 0.028 for all these comparisons, Fig. S5B).

**Figure 4.**
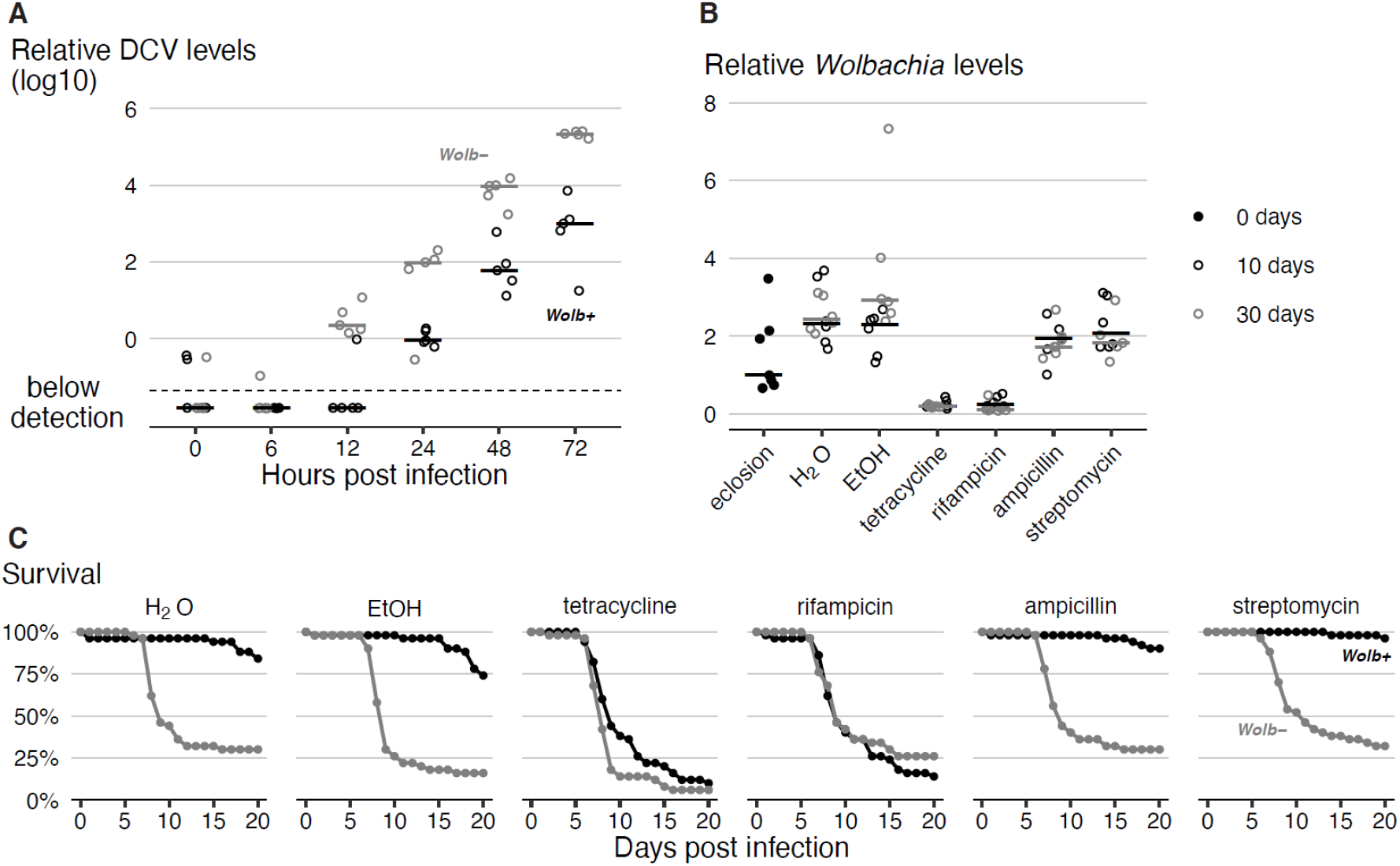
*Wolbachia* presence in adults is required for antiviral protection. (A) Time course analysis of DCV titres in *Wolb+* and *Wolb-* flies raised at 25°C and kept at 18°C after DCV infection. Relative DCV levels were determined by RT-qPCR. Time 0 corresponds to the time of infection. Each dot is a sample consisting of 10 flies, lines are medians. (B) *Wolbachia* levels in single flies raised at 25°C, at eclosion (day 0), after 10 days of different antibiotic and control treatments at 25°C (day 10), and after additional 20 days of treatment at 18°C (day 30), measured by qPCR. A replicate of this experiment is shown in Fig. S5C. (C) Survival of *Wolb+* and *Wolb-* flies, fifty per *Wolbachia* status per treatment, developed at 25°C, collected at eclosion, fed antibiotics-containing food for 10 days at 25°C, infected with DCV (10^8^ TCID_50_/ml), and placed at 18°C on antibiotic-containing food. Mortality was recorded daily. A replicate of this experiment is shown in Fig. S5E.

Developmental temperature determines protection and the protection can be detected as early as we detect viral replication in flies. Thus, we asked if changes in the fly caused by *Wolbachia* during development are sufficient to block viral infection in adults. To answer this, adult flies developed at 25°C were subsequently treated with antibiotics for ten days to remove *Wolbachia* before viral infection (Fig. 4B, S5C). *Wolbachia* levels were reduced approximately 10-fold in ten days of treatment with tetracycline and rifampicin (*p* < 0.001 for both) and stayed low until the end of the experiment at 30 days (*p* < 0.001 for both). These treatments eliminated antiviral protection from the flies (*p* = 1.00 for both, Fig. 4C, S5E). The control treatments with the antibiotics ampicillin and streptomycin did not affect *Wolbachia* levels (*p* > 0.254 for both, Fig. 4B, S5C) and did not affect endosymbiont-mediated protection (in both treatments *Wolbachia* protection is significant, *p* < 0.001 and not different from *Wolbachia* protection in controls, *p* > 0.301, Fig. 4C, S5E). As bacteria in the fly gut were efficiently cleared by all antibiotic treatments (Fig. S5D), we conclude that it is *Wolbachia* loss that leads to the loss of protection and not differential effect of the antibiotics on gut-associated bacteria. These data show that *Wolbachia* presence in adults is required for the *Wolbachia*-conferred antiviral protection, even though the presence or absence of protection is determined by the temperature during development, before the challenge occurs.

### Discussion

Here we show that temperature is a strong modulator of *Wolbachia*-conferred antiviral protection in the natural host *D. melanogaster*. The temperature at which the infection progresses influences this interaction, with *Wolbachia* giving more resistance and increasing survival at lower temperatures. However, the most striking phenotype we report is the effect of host developmental temperature on this interaction. While development at 25°C leads to a strong antiviral protection in terms of survival and resistance to DCV, development at 18°C abrogates, or strongly reduces protection. This is observed with different genotypes of *D. melanogaster*, different variants of *Wolbachia* (*w*Mel and *w*MelCS), and different viruses, and is, therefore, likely to be a general phenomenon.

This complex interaction between temperature and *Wolbachia* protection to viruses may play a role in the natural environment. The outcome of *Wolbachia*-host interactions, in terms of antiviral protection, may vary in regions with different climates or across seasons in the same region. Therefore, *Wolbachia* may behave as a parasite or a mutualist, depending on the conditions. *Wolbachia*’s placement along the parasitism-mutualism continuum at any given time will depend on the cost of harbouring *Wolbachia* and the benefit of antiviral protection to the host. Our results lead to a prediction that *Wolbachia* would protect more in warmer conditions, given the strong reduction in antiviral protection with a low developmental temperature. We have suggested before that the antiviral protection is a fitness advantage that may explain *Wolbachia* prevalence in *D. melanogaster* populations (1). Selection for hosts carrying *Wolbachia* would, therefore, be stronger at higher temperatures. Consequently, geography and seasonality could impact the frequency of *Wolbachia* in a population. Interestingly, there is a high variation in the frequency of *Wolbachia* in *D. melanogaster* natural populations and a clinal distribution of this frequency. At lower latitudes, and therefore warmer climates, the frequency *of Wolbachia* is higher (28). Although, the relationship between *Wolbachia* frequency, temperature and other environmental parameters in *D. melanogaster* is more complex at a smaller geographic scale, *Wolbachia* frequency is the highest in regions with mean annual temperature of 22-26°C (29). In insects, in general, there is also a positive correlation between temperature and *Wolbachia* frequency but only in temperate climates (25).

These results are also relevant for the deployment of *Wolbachia*-carrying mosquitoes to block dengue, Zika, chikungunya and other arboviruses transmission (3, 30–32). It is already known that heat stress impacts *w*Mel in transinfected *Aedes aegypti*, reducing its titres (33), heatwaves can impact titres and frequency of *Wolbachia* in these mosquito populations (34) and at low cycling temperatures *Wolbachia* titers decrease (35). On the other hand, temperatures varying between 25 and 28°C do not seem to affect protection to viruses in this system (18). It would be important, however, to assess how lower temperatures influence *Wolbachia*-induced pathogen blocking in mosquitoes. Determining the temperature range of effectiveness of *Wolbachia*-deploying antiviral field interventions will be crucial to plan where to use it and when to combine it with other complementary approaches (e.g. insecticides).

Pre-infection temperature has a drastic and enigmatic effect on *Wolbachia*-conferred antiviral protection. Lower temperatures during host development determine the level of protection in adult life. Understanding the molecular mechanism of this effect will elucidate how environment interacts with host-microbe symbioses and may be key to understanding *Wolbachia* antiviral protection. The effect of development temperature acts solely on the interaction between virus and *Wolbachia*, and does affect viral infection by itself. The protection conferred by *Wolbachia* seems onset immediately or very soon after viral infection. We observe a difference in viral titres as soon as we detected viral replication in *Wolbachia*-free flies, at 12h after infection. Thus, *Wolbachia* likely inhibits the viral entry or first replication cycle *in vivo. Wolbachia* interference in early stages of viral infection is in agreement with cell culture data (36–38). However, this is not a simple pre-set antiviral state of the host because *Wolbachia* is still required at the time of infection. The lower *Wolbachia* titres in flies developed at 18°C, when compared with flies developed at 25°C, could partly explain a reduction in protection. However, this difference should not be enough for the complete lack of protection when flies develop at 18°C. We have previously observed significant protection to DCV in flies carrying even lower titres of other *w*Mel variants (8). Nonetheless, tissue-specific differences in titres, set during development, may underlie the phenotypic differences. It would, therefore, be interesting to characterize in detail the spatial characterisation of *Wolbachia* tissue tropism and viral infection, in flies developed at the different temperatures. Comparative transcriptomic and metabolomic analysis of *D. melanogaster* and *Wolbachia* in these two conditions could not only elucidate how temperature affects protection, but also the mechanism of *Wolbachia* antiviral protection itself.

Temperature affects many insect-symbiont interactions and their phenotypes (39), including protective symbiosis (40–42). Therefore, this environmental factor may play a general critical role determining the outcome of complex host-endosymbiont-pathogen interactions and shape the geographic distribution of insects and their symbionts. The phenotypic variation we report here, ranging from no protection to strong protection to viruses, indicates that temperature could be a crucial determinant of cost/benefit ratio of carrying *Wolbachia*. Therefore, temperature could deeply impact the *D. melanogaster – Wolbachia* interaction in natural populations. Moreover, these results may have important implications for the deployment of *Wolbachia*-carrying mosquitoes to fight arboviruses transmission and may lead to new approaches to dissect the mechanism of *Wolbachia* antiviral protection.

### Materials and Methods

#### Fly strains and husbandry

DrosDel *w*^1118^ *isogenic D. melanogaster* (*iso*) with *w*MelCS_b *Wolbachia* (*Wolb+*) and the matching control without *Wolbachia* (*Wolb-*) were described elsewhere (1, 8, 43). Aljezur 1 and W-20 *D. melanogaster* lines (*Wolb+* and *Wolb-*) were also described before (1). We determined that the *Wolbachia* variants in both these lines lack an IS5 transposon insertion in gene WD1310, based on the primers described in (44). This insertion is present in all *w*MelCS-like variants, but not in wMel variants (8, 44), therefore Aljezur-1 and W-20 are both *w*Mel-like *Wolbachia* variants. Stocks were maintained at a constant temperature of 25°C on a diet consisting of: 45 g molasses, 75 g sugar, 70 g cornmeal, 20 g yeast extract, 10 g agar, 1100 ml water and 25 ml of 10% Nipagin, with addition of live yeast (Sigma).

#### Virus infection experiments

DCV and FHV were produced and titrated in cell culture as described before (1, 8). Flies for experiments were produced by placing 12 females and 6 males per bottle for 4 days to produce offspring at either 25°C, 18°C or at fluctuating temperature (18°C to 25°C gradual increase during 12 h, and 25°C to 18°C decrease during the subsequent 12 h). After 10 days (25°C), 15 days (fluctuating temperature) or 20 days (18°C) the flies started to eclode. Unless specified otherwise, 0-3 days old flies were collected from the bottles and placed in the vials, ten males per vial, on food without live yeast. Flies were aged for 3 more days at the developmental or otherwise indicated temperature. These 3-6 days old flies were pricked intrathoracically with virus diluted in 50 mM Tris-HCl, pH 7.5. After infection, flies were placed at the indicated temperatures. Survival was monitored daily and vials were changed every 5 days.

#### Nucleic acids extractions and real-time qPCR

DNA for the quantification of *Wolbachia* was extracted from pools of 10 flies using DrosDel protocol (http://www.drosdel.org.uk/molecular_methods.php) (43) or from single flies with the protocol described before (45). RNA for assessment of viral titres was extracted using Trizol (Invitrogen) and cDNA was prepared using M-MLV Reverse Transcriptase (Promega), as described previously (8). Real-time qPCR reactions were carried out in 7900HT Fast Real-Time PCR System (Applied Biosystems) with the iQ™ SYBR® Green supermix (Bio Rad) or in QuantStudio™ 7 Flex Real-Time PCR System (Applied Biosystems) with iTaq™ universal SYBR® Green supermix (Bio Rad). *Wolbachia* was quantified using *wsp* as the target gene and *Drosophila Rpl32* as the reference gene. DCV was quantified with primers for DCV as the target gene and *Drosophila rpl32* as the reference gene. Primers used were: *Wolbachia wsp*, 5’-CATTGGTGTTGGTGTTGGTG-3’ and 5’-ACCGAAATAACGAGCTCCAG-3’; DCV, 5’-TCATCGGTATGCACATTGCT-3’ and 5’-CGCATAACCATGCTCTTCTG-3’; FHV, 5’-ACCTCGATGGCAGGGTTT-3’ and 5’-CTTGAACCATGGCCTTTTG-3’; and *Drosophila rpl32*, 5’-CCGCTTCAAGGGACAGTATC-3’ and 5’-CAATCTCCTTGCGCTTCTTG-3’. Thermal cycling protocol for *Wolbachia* amplification was: initial 50°C for 2 min, denaturation for 10 min at 95°C followed by 40 cycles of 30 s at 95°C, 1 min at 59°C and 30 s at 72°C. For DCV and FHV the same conditions were used, except for an annealing temperature of 56°C. Relative levels of *Wolbachia* or DCV were calculated by the Pfaffl method (46).

#### Antibiotic treatment of flies

*Wolb+* and *Wolb*-flies were raised at 25°C, collected as 0-1 days old adults and placed on fly food with 100 mg/ml of tetracycline hydrochloride, rifampicin, ampicillin sodium or streptomycin sulphate (all Sigma) or control food with antibiotics solvent (water or ethanol). At day 10 of treatment, flies were challenged with DCV, placed at 18°C and survival was followed for additional 20 days. Food vials were changed every three days. At day 0, day 10 and day 30 flies were collected to assay *Wolbachia* levels by qPCR. At day 31, guts from three *Wolb+* flies per condition were dissected to assess the effect of antibiotics on the gut-associated bacteria. Guts were homogenized in 250 µl of sterile LB and 30 µl was plated on mannitol agar plates. This medium sustains growth of the main gut-associated bacteria in lab *D. melanogaster, Acetobacter* and *Lactobacillus* species (47). CFUs were counted after five days of incubation at 26°C.

#### Statistical analysis

All the statistical analysis was performed in R (48). The script of the statistical analysis and output of this analysis are presented in supplementary texts 1 and 2, respectively. All data for statistical analysis and presented in figures are available as supplementary files.

Analysis of survival data was performed with the Cox proportional hazard mixed effect models. Fixed effects, depending on the experiment, included temperature, dose of DCV, presence/absence of *Wolbachia*, and antibiotic treatment, while replicate vials within the same experiment or full experimental replicates were considered as random effects. Model fitting was performed using the coxme package in R version 2.2-16 (49).

*Wolbachia* and DCV titres were analysed with log-transformed qPCR data and linear models or general linear models. Model fitting was performed using lm or the lme4 package in R (50). The effect of interaction between factors in the models was determined by ANOVA. Post hoc analysis of marginal (least square) means were compared between the conditions of interest using the lsmeans package in R (51).

Figures were produced using the ggplot2 package (52).

## Acknowledgments

We thank Dr. Rita Valente for providing assistance with plating bacteria for CFU analyses.

This work was funded by FEBS Long Term Fellowship and Marie Skłodowska-Curie Individual Fellowship MSCA-IF-2017-794507 to EC, FCT grant PTDC/BEX-GMG/3128/2014 to NM, and FCT grants PTDC/BIA-MIC/108327/2008, IF/00839/2015, Wellcome Trust grant 094664/Z/10/Z, and ERC grant 773260 to LT.

---

**Fig. S1.**
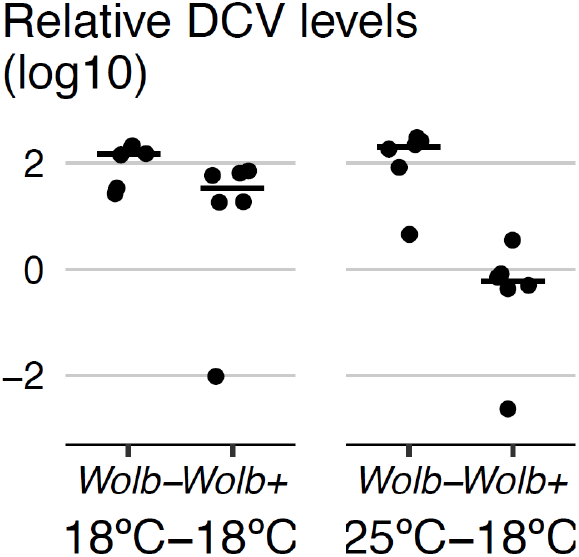
Pre-infection temperature determines viral titres. DCV titres measured by RT-qPCR 3 dpi in *Wolb+* and *Wolb-* flies infected with 10^8^ TCID_50_/ml DCV. Flies were kept at 18°C or 25°C before the infection and at 18°C after the infection. Each dot is a sample, each sample consists of ten flies. Horizontal lines are medians of the replicates. This is a replicate of experiment shown in Fig. 2C.

**Figure S2.**
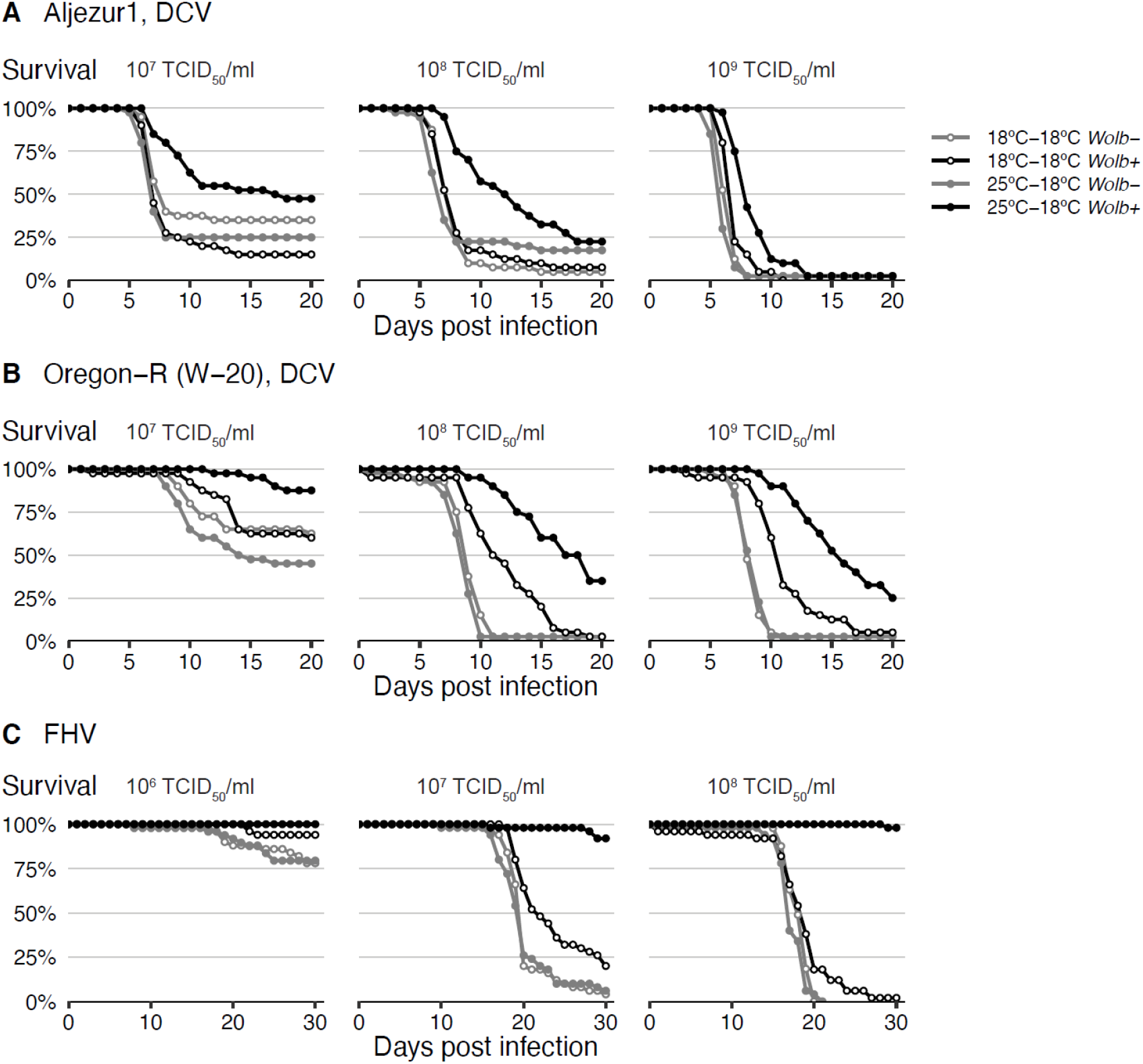
Pre-infection temperature determines *Wolbachia*-conferred antiviral protection in different *Wolbachia* and *Drosophila* genotypes and against different viruses. (A) Aljezur 1 and (B) Oregon-R (W-20) *Wolb+* and *Wolb-* flies, fifty per *Wolbachia* status per condition, were pricked with three dilutions of DCV. (C) *w*^1118^ *iso* flies, fifty per *Wolbachia* status per condition, were infected with three doses of FHV. Flies were kept at 25°C or 18°C before the infection, and at 18°C after the infection. Mortality was recorded daily. This is a replicate of experiment shown in Fig. 3.

**Fig. S3.**
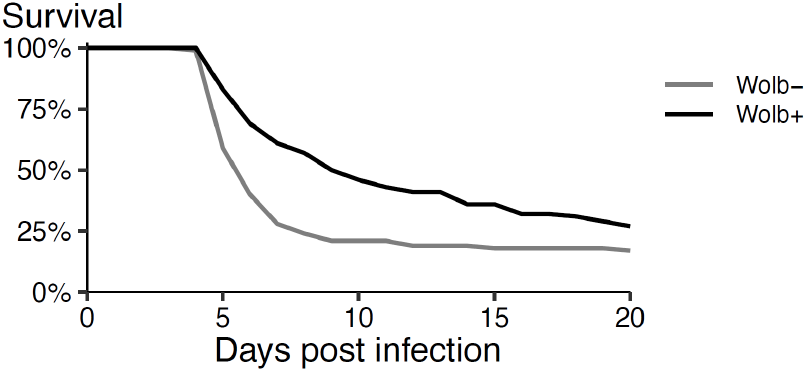
Wolbachia protects flies kept at a temperature cycling daily between 18 and 25°C. Fifty *Wolb+* and *Wolb-* flies were raised with the temperature cycling daily between 25°C (midday) and 18°C (midnight), infected with DCV (10^8^ TCID_50_/ml) and placed at the same cycling temperature after infection. Mortality was recorded daily.

**Figure S4.**
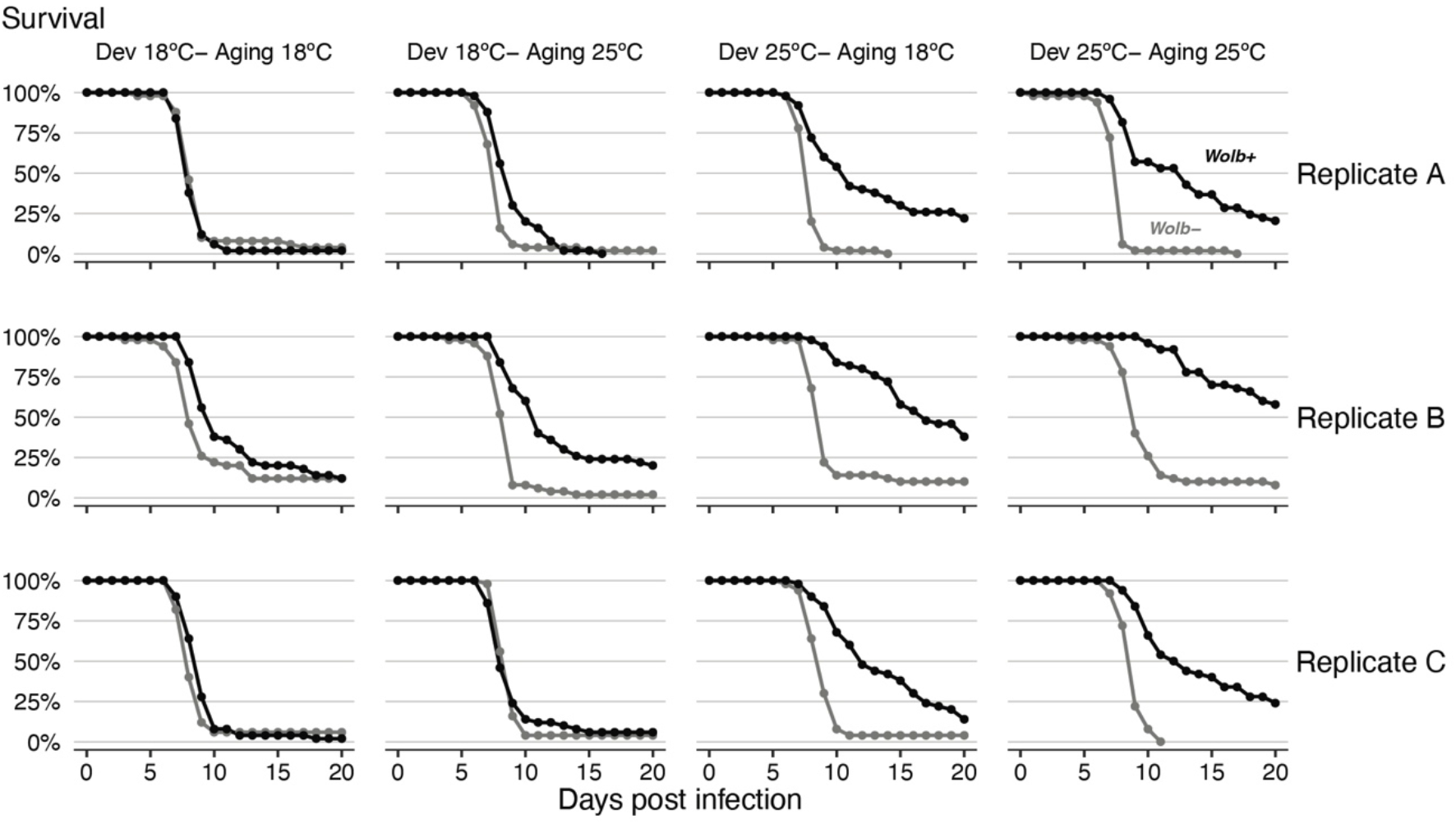
Developmental temperature determines *Wolbachia*-conferred antiviral protection. *Wolb+* and *Wolb-* flies, fifty per *Wolbachia* status per condition per replicate, were raised from egg to adult at 18 or 25°C (Dev), collected at the age of 0-1 days, aged for 3 days at an 18 or 25°C (Aging), pricked with DCV (10^8^ TCID_50_/ml), and placed at 18°C. Mortality was recorded daily. Three replicates of the same experiment are shown.

**Figure S5.**
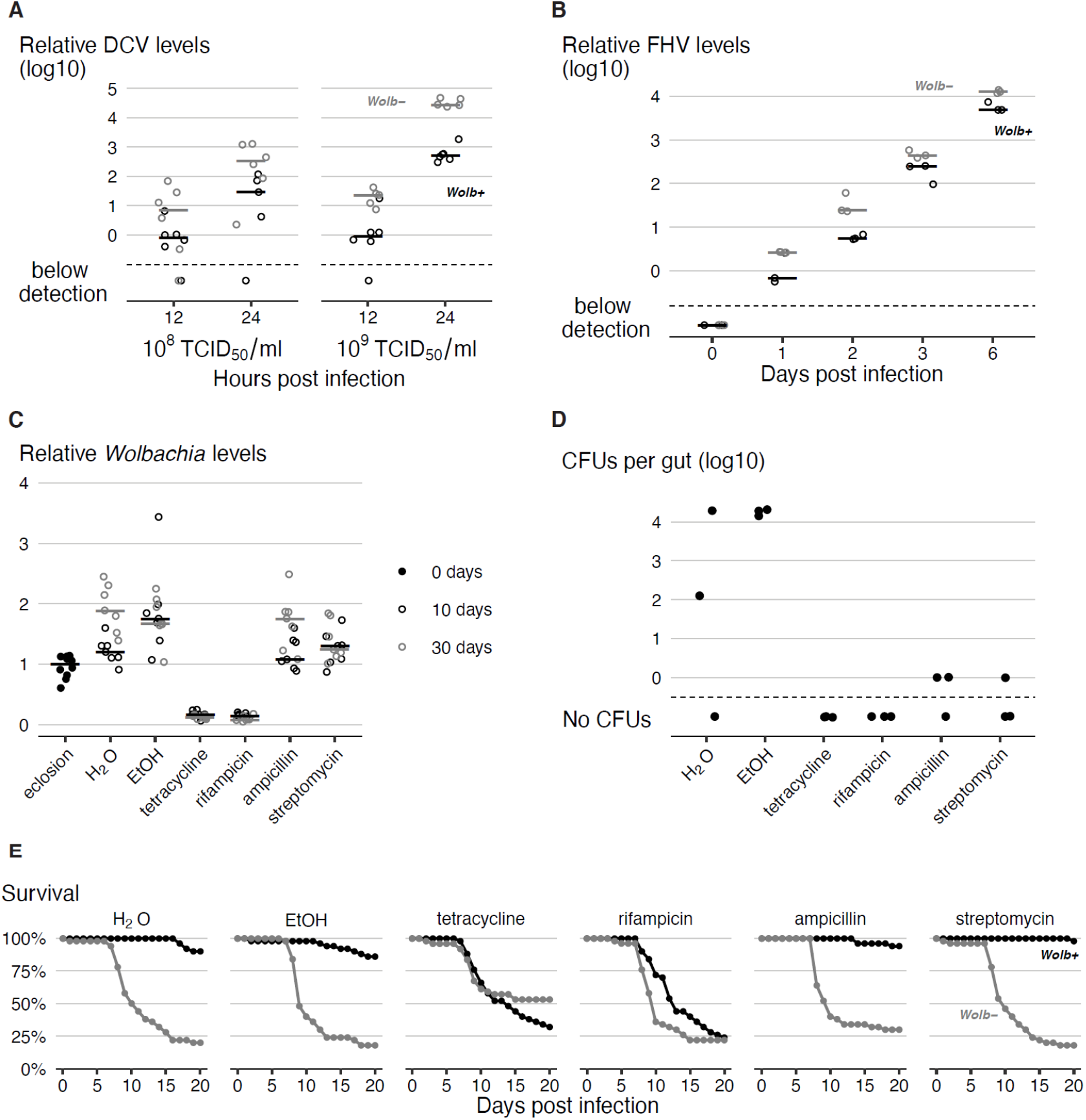
*Wolbachia* presence in adults is required for antiviral protection. (A) DCV titres in *Wolb+* and *Wolb-* flies raised at 25°C and kept at 18°C after DCV infection with two doses, 10^8^ and 10^9^ TCID_50_/ml, and at two time points post infection, 12 and 24h. Relative DCV levels were determined by RT-qPCR. Each dot is a sample consisting of 10 flies, lines are medians. (B) FHV titres in *Wolb+* and *Wolb-* flies raised at 25°C and kept at 25°C after infection. Each dot is a sample consisting of 10 flies, lines are medians, and time 0 corresponds to the time of infection. (C) *Wolbachia* levels in single flies raised at 25°C, at eclosion (day 0), after 10 days of different antibiotic/ control treatments at 25°C (day 10), and after additional 20 days of treatment at 18°C (day 30) measured by qPCR. This is the replicate of the experiment shown in Fig. 4B. (D) Colony forming units (CFUs) in dissected *Wolb+* fly guts at the day 30 of each antibiotic or control treatment. Each dot is a pool of three guts. (E) Survival of *Wolb+* and *Wolb-* flies, fifty per *Wolbachia* status per treatment, developed at 25°C, collected at eclosion, fed antibiotics-containing food for 10 days at 25°C, infected with DCV (10^8^ TCID_50_/ml), and placed at 18°C on antibiotic-containing food. Mortality was recorded daily. This is the replicate of the experiment shown in Fig. 4C.

## Supplementary Information

**SI Text S1 (text_s1**.**txt)**

Rmd script for statistical analysis

**SI Text S2 (text_s2**.**pdf)**

Output of statistical analysis

**SI Dataset S1 (dataset_s1**.**txt)**

Data for figures 1A, B

**SI Dataset S2 (dataset_s2**.**txt)**

Data for figure 1C

**SI Dataset S3 (dataset_s3**.**txt)**

Data for figure 1D

**SI Dataset S4 (dataset_s4**.**txt)**

Data for figures 2A, B, and S3

**SI Dataset S5 (dataset_s5**.**txt)**

Data for figures 2C and S1

**SI Dataset S6 (dataset_s6**.**txt)**

Data for figure 2D

**SI Dataset S7 (dataset_s7**.**txt)**

Data for figures 3A, B, and S2A, B

**SI Dataset S8 (dataset_s8**.**txt)**

Data for figures 3C and S2C

**SI Dataset S9 (dataset_s9**.**txt)**

Data for figure S4

**SI Dataset S10 (dataset_s10**.**txt)**

Data for figure 4A

**SI Dataset S11 (dataset_s11**.**txt)**

Data for figure S5A

**SI Dataset S12 (dataset_s12**.**txt)**

Data for figure S5B

**SI Dataset S13 (dataset_s13**.**txt)**

Data for figures 4B and S5C

**SI Dataset S14 (dataset_s14**.**txt)**

Data for figures 4C and S5E

**SI Dataset S15 (dataset_s15**.**txt)**

Data for figure S5D

