## Supplementary material for "*Wolbachia-conferred* antiviral protection is determined by developmental temperature": Text S2

### Statistical analysis of temperature effect on *Wolbachia*-induced protection to viruses

24 June, 2020

**Session info** **49**

### Figure 1

#### Survival at different post-infections temperatures

```
surv_2temp<-fread("dataset_s1.txt")

#make a variable describing each unique set of conditions
surv_2temp[,RepFull:=interaction(Temp,Wolb,Dose,Replicate,sep = "_")]
surv_2temp<-surv_2temp[,lapply(.SD,char_asfactor)]

##Diagnostics
```

```r
#How many individuals were tested
ftable(xtabs(~Temp+Dose+Wolb,surv_2temp))
```

```
##           Wolb Wolb- Wolb+
## Temp Dose
## 18C  E5          50    50
##      E6          50    50
##      E7          50    50
##      E8          50    50
##      E9          50    50
## 25C  E5          50    50
##      E6          50    50
##      E7          50    50
##      E8          50    50
##      E9          50    50
```

```r
##Data analysis
# Wolbachia * Dose * Temperature comparisons
## Full model

cox_2temp_full<-coxme(Surv(Time,Status)~Wolb*Dose*Temp+(1|RepFull), surv_2temp)

# Anova Table
Anova(cox_2temp_full,test.statistic = "LR")
```

```
## Analysis of Deviance Table (Type II tests)
##                LR Chisq Df Pr(>Chisq)
## Wolb             63.810  1  1.370e-15 ***
## Dose            220.370  4  < 2.2e-16 ***
## Temp             98.543  1  < 2.2e-16 ***
## Wolb:Dose         8.001  4    0.09154 .
## Wolb:Temp        19.497  1  1.007e-05 ***
## Dose:Temp        30.601  4  3.693e-06 ***
## Wolb:Dose:Temp    9.123  4    0.05810 .
## ---
## Signif. codes:  0 '***' 0.001 '**' 0.01 '*' 0.05 '.' 0.1 ' ' 1
```

```r
# minimum model

cox_2temp_final<-coxme(Surv(Time,Status)~Temp*(Dose+Wolb)+(1|RepFull), surv_2temp)

Anova(cox_2temp_final,test.statistic = "LR")
```

```
## Analysis of Deviance Table (Type II tests)
##           LR Chisq Df Pr(>Chisq)
## Temp        94.853  1  < 2.2e-16 ***
## Dose       220.370  4  < 2.2e-16 ***
## Wolb        63.810  1  1.370e-15 ***
## Temp:Dose   31.862  4  2.041e-06 ***
## Temp:Wolb   17.551  1  2.798e-05 ***
## ---
## Signif. codes:  0 '***' 0.001 '**' 0.01 '*' 0.05 '.' 0.1 ' ' 1
```

```r
summary(cox_2temp_final)
```

```
## Cox mixed-effects model fit by maximum likelihood
```

```
##   Data: surv_2temp
##   events, n = 655, 1000
##   Iterations= 17 126
##                     NULL Integrated    Fitted
## Log-likelihood -4237.264  -3581.165 -3523.542
##
##                     Chisq    df p     AIC     BIC
## Integrated loglik 1312.20 12.00 0 1288.20 1234.38
##  Penalized loglik 1427.44 52.22 0 1323.01 1088.85
##
## Model:  Surv(Time, Status) ~ Temp * (Dose + Wolb) + (1 | RepFull)
## Fixed coefficients
##                         coef  exp(coef)  se(coef)     z       p
## Temp25C           -0.3599373  0.6977201 0.5636204 -0.64 5.2e-01
## DoseE6             1.6146846  5.0263023 0.4537914  3.56 3.7e-04
## DoseE7             3.3818780 29.4259800 0.4337873  7.80 6.3e-15
## DoseE8             3.8383442 46.4485027 0.4323228  8.88 0.0e+00
## DoseE9             4.1150815 61.2572096 0.4318331  9.53 0.0e+00
## WolbWolb+         -1.8410024  0.1586583 0.1885979 -9.76 0.0e+00
## Temp25C:DoseE6     0.9830450  2.6725819 0.6366543  1.54 1.2e-01
## Temp25C:DoseE7     2.2328572  9.3264756 0.6158195  3.63 2.9e-04
## Temp25C:DoseE8     2.8684844 17.6103079 0.6146590  4.67 3.1e-06
## Temp25C:DoseE9     2.3895115 10.9081641 0.6126688  3.90 9.6e-05
## Temp25C:WolbWolb+  1.0934571  2.9845742 0.2510025  4.36 1.3e-05
##
## Random effects
##  Group   Variable  Std Dev   Variance
##  RepFull Intercept 0.4178493 0.1745981
```

Significant interaction between *Wolbachia* and Temperature, and between Dose and Temperature. No interaction between *Wolbachia* and Dose.

```
#Comparisons between hazard ratios

#Hazards ratios between wolb+ and wolb- at both temp
contr_2temp_Wolb<-lsmeans::lsmeans(cox_2temp_final,pairwise~Wolb|Temp)
contr_2temp_Wolb
```

```
## $lsmeans
## Temp = 18C:
##  Wolb   lsmean    SE  df asymp.LCL asymp.UCL
##  Wolb- -0.0203 0.109 Inf    -0.234     0.194
##  Wolb+ -1.8613 0.137 Inf    -2.129    -1.593
##
## Temp = 25C:
##  Wolb   lsmean    SE  df asymp.LCL asymp.UCL
##  Wolb-  1.3146 0.116 Inf     1.087     1.542
##  Wolb+  0.5670 0.118 Inf     0.335     0.799
##
## Results are averaged over the levels of: Dose
## Results are given on the log (not the response) scale.
## Confidence level used: 0.95
##
```

```
## $contrasts
## Temp = 18C:
##  contrast      estimate    SE  df z.ratio p.value
##  Wolb- - Wolb+    1.841 0.189 Inf 9.762   <.0001
##
## Temp = 25C:
##  contrast      estimate    SE  df z.ratio p.value
##  Wolb- - Wolb+    0.748 0.171 Inf 4.372   <.0001
##
## Results are averaged over the levels of: Dose
## Results are given on the log (not the response) scale.
```

```r
contrast(contr_2temp_Wolb$contrasts,method="pairwise",by="contrast")
```

```
## contrast = Wolb- - Wolb+:
##  contrast1 estimate    SE  df z.ratio p.value
##  18C - 25C     1.09 0.251 Inf 4.356   <.0001
##
## Results are averaged over the levels of: Dose
## Results are given on the log (not the response) scale.
```

*Wolbachia* has an effect at both temperatures, significantly stronger effect at 18ºC post-infection temperature.

```r
#Hazards ratios between temperatures, at all doses
mcp_temps_final<-lsmeans::lsmeans(cox_2temp_final,pairwise~Temp|Dose)
mcp_temps_final
```

```
## $lsmeans
## Dose = E5:
##  Temp lsmean    SE  df asymp.LCL asymp.UCL
##  18C  -3.531 0.364 Inf   -4.2434    -2.818
##  25C  -3.344 0.371 Inf   -4.0711    -2.617
##
## Dose = E6:
##  Temp lsmean    SE  df asymp.LCL asymp.UCL
##  18C  -1.916 0.229 Inf   -2.3655    -1.467
##  25C  -0.746 0.192 Inf   -1.1222    -0.370
##
## Dose = E7:
##  Temp lsmean    SE  df asymp.LCL asymp.UCL
##  18C  -0.149 0.179 Inf   -0.4990     0.201
##  25C   2.271 0.176 Inf    1.9256     2.616
##
## Dose = E8:
##  Temp lsmean    SE  df asymp.LCL asymp.UCL
##  18C   0.308 0.172 Inf   -0.0299     0.645
##  25C   3.363 0.185 Inf    3.0010     3.725
##
## Dose = E9:
##  Temp lsmean    SE  df asymp.LCL asymp.UCL
##  18C   0.584 0.169 Inf    0.2526     0.916
##  25C   3.161 0.181 Inf    2.8050     3.516
##
```

```
## Results are averaged over the levels of: Wolb
## Results are given on the log (not the response) scale.
## Confidence level used: 0.95
##
## $contrasts
## Dose = E5:
##  contrast  estimate    SE  df z.ratio p.value
##  18C - 25C   -0.187 0.563 Inf  -0.332 0.7402
##
## Dose = E6:
##  contrast  estimate    SE  df z.ratio p.value
##  18C - 25C   -1.170 0.308 Inf  -3.804 0.0001
##
## Dose = E7:
##  contrast  estimate    SE  df z.ratio p.value
##  18C - 25C   -2.420 0.254 Inf  -9.527 <.0001
##
## Dose = E8:
##  contrast  estimate    SE  df z.ratio p.value
##  18C - 25C   -3.055 0.251 Inf -12.170 <.0001
##
## Dose = E9:
##  contrast  estimate    SE  df z.ratio p.value
##  18C - 25C   -2.576 0.245 Inf -10.514 <.0001
##
## Results are averaged over the levels of: Wolb
## Results are given on the log (not the response) scale.
```

Temperature has a significant effect in survival, except for lower dose ($10^5$ TCID/$_{50}$), which has very low lethality even without *Wolbachia*.

**Figures 1A and B**

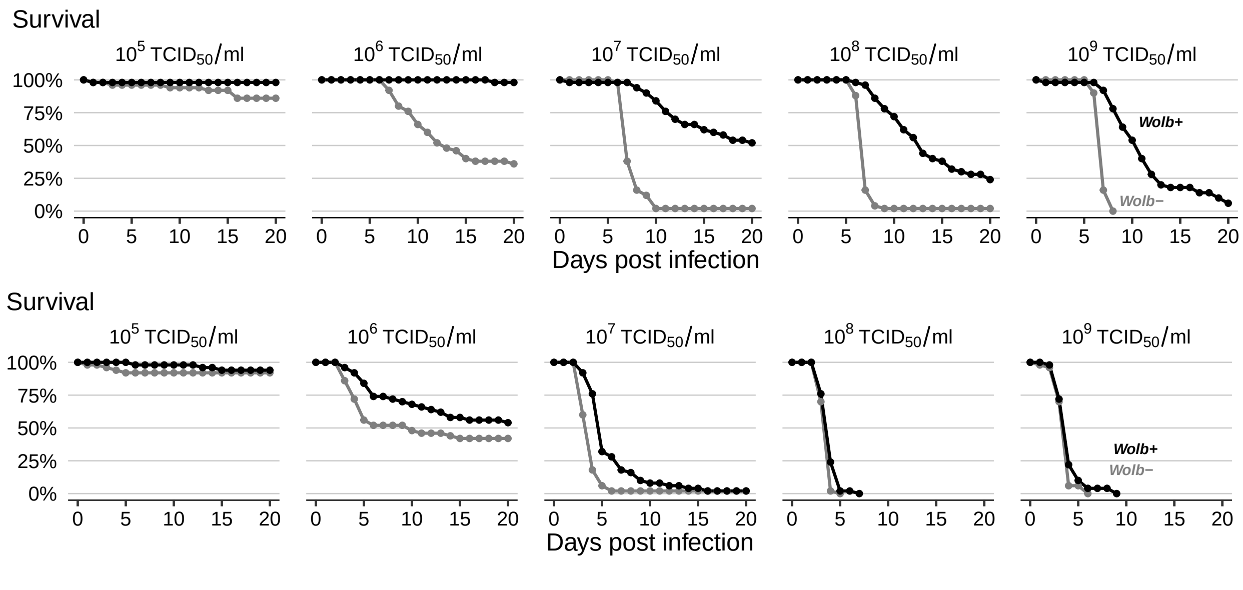

#### DCV Titers at different post-infections temperatures

```r
#Read the data
DCV_bytemp<-fread("dataset_s2.txt")
DCV_bytemp<-DCV_bytemp[,lapply(.SD,char_asfactor)]
DCV_bytemp[,logRatio:=log10(Ratio)][,inter:=interaction(Wolb,Temp,Dose)]
str(DCV_bytemp)
```

```
## Classes 'data.table' and 'data.frame':   100 obs. of  7 variables:
##  $ Sample  : Factor w/ 2 levels "138","iso": 1 1 1 1 1 1 1 1 1 1 ...
##  $ Wolb    : Factor w/ 2 levels "Wolb-","Wolb+": 2 2 2 2 2 2 2 2 2 2 ...
##  $ Ratio   : num  6.06e-04 2.77e-05 3.21e-05 9.63e-06 2.28e-05 ...
##  $ Temp    : Factor w/ 2 levels "18C","25C": 1 1 1 1 1 1 1 1 1 1 ...
##  $ Dose    : Factor w/ 5 levels "E5","E6","E7",..: 1 1 1 1 1 2 2 2 2 2 ...
##  $ logRatio: num  -3.22 -4.56 -4.49 -5.02 -4.64 ...
##  $ inter   : Factor w/ 20 levels "Wolb-.18C.E5",..: 2 2 2 2 2 6 6 6 6 6 ...
##  - attr(*, ".internal.selfref")=<externalptr>
```

```r
##Diagnostics
#How many individuals were tested
ftable(xtabs(~Wolb+Temp+Dose,data=DCV_bytemp))
```

```
##             Dose E5 E6 E7 E8 E9
## Wolb  Temp
## Wolb- 18C         5  5  5  5  5
##       25C         5  5  5  5  5
## Wolb+ 18C         5  5  5  5  5
##       25C         5  5  5  5  5
```

```r
##Data analysis
#linear model
mod.log<-lm(logRatio~Wolb*Temp*Dose,data=DCV_bytemp)
Anova(mod.log)
```

```
## Anova Table (Type II tests)
##
## Response: logRatio
##                 Sum Sq Df  F value    Pr(>F)
## Wolb           122.434  1 164.4990 < 2.2e-16 ***
## Temp           112.567  1 151.2427 < 2.2e-16 ***
## Dose           107.331  4  36.0517 < 2.2e-16 ***
## Wolb:Temp        6.928  1   9.3086  0.003094 **
## Wolb:Dose        3.087  4   1.0369  0.393453
## Temp:Dose        4.650  4   1.5620  0.192507
## Wolb:Temp:Dose  24.034  4   8.0729  1.61e-05 ***
## Residuals       59.543 80
## ---
## Signif. codes:  0 '***' 0.001 '**' 0.01 '*' 0.05 '.' 0.1 ' ' 1
```

```r
summary(mod.log)
```

```
##
## Call:
## lm(formula = logRatio ~ Wolb * Temp * Dose, data = DCV_bytemp)
##
## Residuals:
##     Min      1Q  Median      3Q     Max
```

```
## -2.8369 -0.2048  0.0012  0.1655  3.1067
##
## Coefficients:
##                            Estimate Std. Error t value Pr(>|t|)
## (Intercept)                 -3.7890     0.3858  -9.821 2.16e-15 ***
## WolbWolb+                   -0.5964     0.5456  -1.093  0.27762
## Temp25C                      3.1187     0.5456   5.716 1.81e-07 ***
## DoseE6                       1.2352     0.5456   2.264  0.02629 *
## DoseE7                       3.5966     0.5456   6.592 4.27e-09 ***
## DoseE8                       4.0454     0.5456   7.414 1.12e-10 ***
## DoseE9                       4.3099     0.5456   7.899 1.28e-11 ***
## WolbWolb+:Temp25C           -2.2382     0.7716  -2.901  0.00481 **
## WolbWolb+:DoseE6            -1.5190     0.7716  -1.969  0.05247 .
## WolbWolb+:DoseE7            -3.2261     0.7716  -4.181 7.38e-05 ***
## WolbWolb+:DoseE8            -3.4387     0.7716  -4.456 2.68e-05 ***
## WolbWolb+:DoseE9            -2.5312     0.7716  -3.280  0.00154 **
## Temp25C:DoseE6              -0.2890     0.7716  -0.375  0.70899
## Temp25C:DoseE7              -2.0023     0.7716  -2.595  0.01125 *
## Temp25C:DoseE8              -2.4612     0.7716  -3.190  0.00203 **
## Temp25C:DoseE9              -2.8632     0.7716  -3.711  0.00038 ***
## WolbWolb+:Temp25C:DoseE6     2.0791     1.0913   1.905  0.06035 .
## WolbWolb+:Temp25C:DoseE7     4.6631     1.0913   4.273 5.28e-05 ***
## WolbWolb+:Temp25C:DoseE8     5.1238     1.0913   4.695 1.09e-05 ***
## WolbWolb+:Temp25C:DoseE9     4.5895     1.0913   4.206 6.74e-05 ***
## ---
## Signif. codes:  0 '***' 0.001 '**' 0.01 '*' 0.05 '.' 0.1 ' ' 1
##
## Residual standard error: 0.8627 on 80 degrees of freedom
## Multiple R-squared:  0.8649, Adjusted R-squared:  0.8328
## F-statistic: 26.94 on 19 and 80 DF,  p-value: < 2.2e-16
```

Interaction between *Wolbachia*, Dose, and Temperature

```
#Least square means

#effect of Wolbachia at different doses, at different temperatures
lsm_bywolb.log<-lsmeans::lsmeans(mod.log,pairwise~Wolb|Dose:Temp,adj="none")
summary(lsm_bywolb.log,by=NULL,adj="holm")
```

```
## $lsmeans
##  Wolb  Dose Temp lsmean    SE df lower.CL upper.CL
##  Wolb- E5   18C  -3.789 0.386 80   -4.994   -2.584
##  Wolb+ E5   18C  -4.385 0.386 80   -5.590   -3.181
##  Wolb- E6   18C  -2.554 0.386 80   -3.758   -1.349
##  Wolb+ E6   18C  -4.669 0.386 80   -5.874   -3.465
##  Wolb- E7   18C  -0.192 0.386 80   -1.397    1.012
##  Wolb+ E7   18C  -4.015 0.386 80   -5.219   -2.810
##  Wolb- E8   18C   0.256 0.386 80   -0.948    1.461
##  Wolb+ E8   18C  -3.779 0.386 80   -4.983   -2.574
##  Wolb- E9   18C   0.521 0.386 80   -0.684    1.725
##  Wolb+ E9   18C  -2.607 0.386 80   -3.811   -1.402
##  Wolb- E5   25C  -0.670 0.386 80   -1.875    0.534
##  Wolb+ E5   25C  -3.505 0.386 80   -4.710   -2.300
```

```
##  Wolb- E6   25C   0.276 0.386 80   -0.929    1.480
##  Wolb+ E6   25C  -1.999 0.386 80   -3.203   -0.794
##  Wolb- E7   25C   0.924 0.386 80   -0.281    2.128
##  Wolb+ E7   25C  -0.474 0.386 80   -1.678    0.731
##  Wolb- E8   25C   0.914 0.386 80   -0.291    2.118
##  Wolb+ E8   25C  -0.236 0.386 80   -1.440    0.969
##  Wolb- E9   25C   0.776 0.386 80   -0.428    1.981
##  Wolb+ E9   25C   0.000 0.386 80   -1.205    1.205
##
## Confidence level used: 0.95
## Conf-level adjustment: bonferroni method for 20 estimates
##
## $contrasts
##  contrast      Dose Temp estimate    SE df t.ratio p.value
##  Wolb- - Wolb+ E5   18C     0.596 0.546 80 1.093   0.3173
##  Wolb- - Wolb+ E6   18C     2.115 0.546 80 3.877   0.0011
##  Wolb- - Wolb+ E7   18C     3.822 0.546 80 7.006   <.0001
##  Wolb- - Wolb+ E8   18C     4.035 0.546 80 7.395   <.0001
##  Wolb- - Wolb+ E9   18C     3.128 0.546 80 5.732   <.0001
##  Wolb- - Wolb+ E5   25C     2.835 0.546 80 5.195   <.0001
##  Wolb- - Wolb+ E6   25C     2.275 0.546 80 4.169   0.0005
##  Wolb- - Wolb+ E7   25C     1.398 0.546 80 2.561   0.0492
##  Wolb- - Wolb+ E8   25C     1.150 0.546 80 2.107   0.1148
##  Wolb- - Wolb+ E9   25C     0.776 0.546 80 1.423   0.3173
##
## P value adjustment: holm method for 10 tests
```

*Wolbachia* has an effect at most doses and temperatures combinations. At 18ºC *Wolbachia* effect is significant at all doses except lowest one. At 25ºC *Wolbachia* effect is significant at the three lowest doses (out of 5) doses.

```r
#effect of Wolbachia at different temperatures
lsm_bywolbtemp.log<-lsmeans::lsmeans(mod.log,pairwise~Wolb|Temp,adj="none")
```

```
## NOTE: Results may be misleading due to involvement in interactions
```

```r
summary(lsm_bywolbtemp.log,by=NULL,adj="holm")
```

```
## $lsmeans
##  Wolb  Temp lsmean    SE df lower.CL upper.CL
##  Wolb- 18C  -1.152 0.173 80 -1.59250   -0.711
##  Wolb+ 18C  -3.891 0.173 80 -4.33193   -3.450
##  Wolb- 25C   0.444 0.173 80  0.00302    0.885
##  Wolb+ 25C  -1.243 0.173 80 -1.68354   -0.802
##
## Results are averaged over the levels of: Dose
## Confidence level used: 0.95
## Conf-level adjustment: bonferroni method for 4 estimates
##
## $contrasts
##  contrast      Temp estimate    SE df t.ratio p.value
##  Wolb- - Wolb+ 18C      2.74 0.244 80 11.227  <.0001
##  Wolb- - Wolb+ 25C      1.69 0.244 80  6.912  <.0001
##
```

```
## Results are averaged over the levels of: Dose
## P value adjustment: holm method for 2 tests
```

```
#fold difference 18C
log_bywolbtemp_18C <- summary(lsm_bywolbtemp.log,by=NULL,adj="holm")$contrasts[1,3]
'^'(10,log_bywolbtemp_18C)
```

```
## [1] 548.8167
```

```
#fold difference 25C
log_bywolbtemp_25C <- summary(lsm_bywolbtemp.log,by=NULL,adj="holm")$contrasts[2,3]
'^'(10,log_bywolbtemp_25C)
```

```
## [1] 48.5923
```

```
#Directly test the differences
contrast(lsm_bywolbtemp.log$contrasts,"pairwise",by="contrast")
```

```
## contrast = Wolb- - Wolb+:
##  contrast1 estimate    SE df t.ratio p.value
##  18C - 25C     1.05 0.345 80 3.051   0.0031
##
## Results are averaged over the levels of: Dose
```

On average *Wolbachia* gives more resistance at 18ºC

```
#effect of temp at different doses and Wolb presence
lsm_byTemp.log<-lsmeans::lsmeans(mod.log,pairwise~Temp|Dose:Wolb,adj="none")
summary(lsm_byTemp.log,by=NULL,adj="holm")
```

```
## $lsmeans
##  Temp Dose Wolb  lsmean    SE df lower.CL upper.CL
##  18C  E5   Wolb- -3.789 0.386 80   -4.994   -2.584
##  25C  E5   Wolb- -0.670 0.386 80   -1.875    0.534
##  18C  E6   Wolb- -2.554 0.386 80   -3.758   -1.349
##  25C  E6   Wolb-  0.276 0.386 80   -0.929    1.480
##  18C  E7   Wolb- -0.192 0.386 80   -1.397    1.012
##  25C  E7   Wolb-  0.924 0.386 80   -0.281    2.128
##  18C  E8   Wolb-  0.256 0.386 80   -0.948    1.461
##  25C  E8   Wolb-  0.914 0.386 80   -0.291    2.118
##  18C  E9   Wolb-  0.521 0.386 80   -0.684    1.725
##  25C  E9   Wolb-  0.776 0.386 80   -0.428    1.981
##  18C  E5   Wolb+ -4.385 0.386 80   -5.590   -3.181
##  25C  E5   Wolb+ -3.505 0.386 80   -4.710   -2.300
##  18C  E6   Wolb+ -4.669 0.386 80   -5.874   -3.465
##  25C  E6   Wolb+ -1.999 0.386 80   -3.203   -0.794
##  18C  E7   Wolb+ -4.015 0.386 80   -5.219   -2.810
##  25C  E7   Wolb+ -0.474 0.386 80   -1.678    0.731
##  18C  E8   Wolb+ -3.779 0.386 80   -4.983   -2.574
##  25C  E8   Wolb+ -0.236 0.386 80   -1.440    0.969
##  18C  E9   Wolb+ -2.607 0.386 80   -3.811   -1.402
##  25C  E9   Wolb+  0.000 0.386 80   -1.205    1.205
##
## Confidence level used: 0.95
## Conf-level adjustment: bonferroni method for 20 estimates
```

```
##
## $contrasts
##  contrast  Dose Wolb  estimate    SE df t.ratio p.value
##  18C - 25C E5   Wolb-   -3.119 0.546 80 -5.716  <.0001
##  18C - 25C E6   Wolb-   -2.830 0.546 80 -5.186  <.0001
##  18C - 25C E7   Wolb-   -1.116 0.546 80 -2.046  0.1762
##  18C - 25C E8   Wolb-   -0.657 0.546 80 -1.205  0.4636
##  18C - 25C E9   Wolb-   -0.255 0.546 80 -0.468  0.6409
##  18C - 25C E5   Wolb+   -0.880 0.546 80 -1.614  0.3317
##  18C - 25C E6   Wolb+   -2.671 0.546 80 -4.894  <.0001
##  18C - 25C E7   Wolb+   -3.541 0.546 80 -6.490  <.0001
##  18C - 25C E8   Wolb+   -3.543 0.546 80 -6.493  <.0001
##  18C - 25C E9   Wolb+   -2.607 0.546 80 -4.777  <.0001
##
## P value adjustment: holm method for 10 tests
```

```
#average effect of temperature
lsm_byTemponly.log<-lsmeans::lsmeans(mod.log,pairwise~Temp,adj="none")
```

```
## NOTE: Results may be misleading due to involvement in interactions
```

```
summary(lsm_byTemponly.log,by=NULL,adj="holm")
```

```
## $lsmeans
##  Temp lsmean    SE df lower.CL upper.CL
##  18C  -2.521 0.122 80   -2.800   -2.243
##  25C  -0.399 0.122 80   -0.678   -0.121
##
## Results are averaged over the levels of: Wolb, Dose
## Confidence level used: 0.95
## Conf-level adjustment: bonferroni method for 2 estimates
##
## $contrasts
##  contrast  estimate    SE df t.ratio p.value
##  18C - 25C    -2.12 0.173 80 -12.298 <.0001
##
## Results are averaged over the levels of: Wolb, Dose
```

On average DCV titres are lower at 18ºC than 25ºC

```
#average effect of Wolbachia
lsm_byWolbonly.log<-lsmeans::lsmeans(mod.log,pairwise~Wolb,adj="none")
```

```
## NOTE: Results may be misleading due to involvement in interactions
```

```
summary(lsm_byWolbonly.log,by=NULL,adj="holm")
```

```
## $lsmeans
##  Wolb   lsmean    SE df lower.CL upper.CL
##  Wolb- -0.354 0.122 80   -0.633  -0.0751
##  Wolb+ -2.567 0.122 80   -2.846  -2.2881
##
## Results are averaged over the levels of: Temp, Dose
## Confidence level used: 0.95
## Conf-level adjustment: bonferroni method for 2 estimates
```

```
##
## $contrasts
##  contrast      estimate    SE df t.ratio p.value
##  Wolb- - Wolb+     2.21 0.173 80 12.826  <.0001
##
## Results are averaged over the levels of: Temp, Dose
```

##### Figure 1C

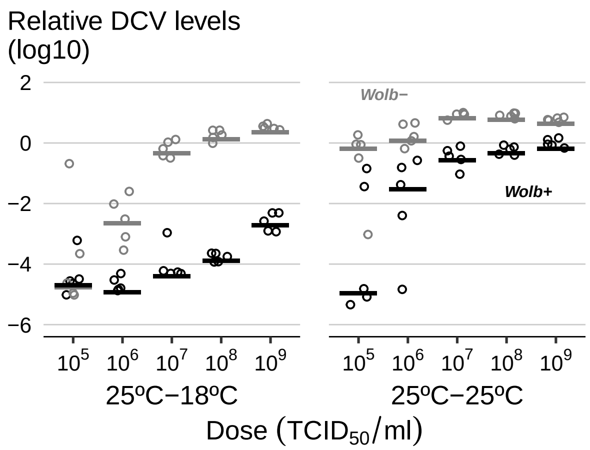

#### *Wolbachia* levels post-infection at different temperatures

```
wolb_DCV<-fread("dataset_s3.txt")
unique(wolb_DCV$Treatment)
```

```
## [1] "buffer" "CTRL"   "DCV"
```

```
wolb_DCV[,pre_post:=interaction(Pre_Temp,Post_Temp,sep="_")][,inter:=droplevels(interaction(Treatment,Da
wolb_DCV[,logWolb:=log10(Wolb)][,inter:=relevel(inter,"CTRL.day3.25C_18C")]
wolb_DCV<-wolb_DCV[,lapply(.SD,char_asfactor)]
#remove day 0 from analysis
wolb_DCV_day3 <- filter(wolb_DCV, Day != "day0")
head(wolb_DCV)
```

```
##                Sample Treatment  Day Pre_Temp Post_Temp      Wolb pre_post
## 1: buffer_day_3_18C_a    buffer day3      25C       18C 1.4373219  25C_18C
## 2: buffer_day_3_18C_b    buffer day3      25C       18C 1.5426671  25C_18C
## 3: buffer_day_3_18C_c    buffer day3      25C       18C 1.5763952  25C_18C
## 4: buffer_day_3_18C_d    buffer day3      25C       18C 1.3177769  25C_18C
## 5: buffer_day_3_18C_e    buffer day3      25C       18C 1.0030623  25C_18C
## 6: buffer_day_3_25C_a    buffer day3      25C       25C 0.9432642  25C_25C
##                inter      logWolb
## 1: buffer.day3.25C_18C  0.157554047
## 2: buffer.day3.25C_18C  0.188272227
## 3: buffer.day3.25C_18C  0.197665103
## 4: buffer.day3.25C_18C  0.119841880
## 5: buffer.day3.25C_18C  0.001327906
## 6: buffer.day3.25C_25C -0.025366649
```

```r
#lm
wolb_model<-lm(logWolb~Treatment*Post_Temp,data=wolb_DCV_day3)
Anova(wolb_model)
```

```
## Anova Table (Type II tests)
##
## Response: logWolb
##                        Sum Sq Df F value Pr(>F)
## Treatment            0.016584  2  0.8847 0.4259
## Post_Temp            0.000217  1  0.0232 0.8803
## Treatment:Post_Temp  0.013996  2  0.7466 0.4846
## Residuals            0.224945 24
```

No effect of DCV infection or post-infection temperature on *Wolbachia* levels

##### Figure 1D

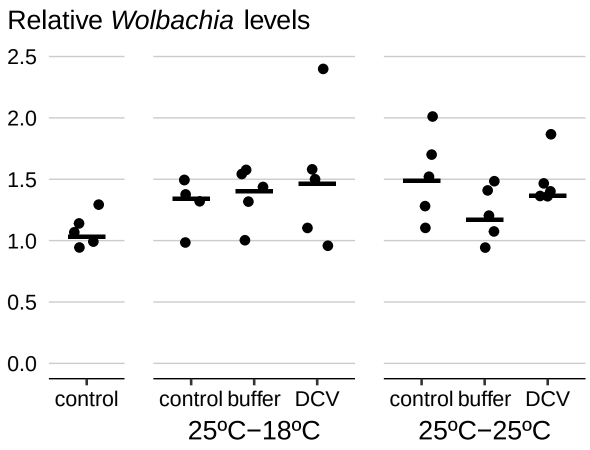

### Figure 2 and S1

#### Effect of different pre and post-infection temperatures on survival

```r
##Data import and create column with replicate information
surv_pre_post_ini<-fread("dataset_s4.txt")

surv_pre_post_ini<-surv_pre_post_ini[,":="(Pre_Temp=as.factor(Pre_Temp),
                    Post_Temp=as.factor(Post_Temp),
                    RepFull=interaction(Wolb,Pre_Temp,Post_Temp,Replicate,sep = "_"))
                    ][,lapply(.SD,char_asfactor)]

surv_cyc<-droplevels(surv_pre_post_ini[Pre_Temp=="cycling"])
surv_pre_post<-droplevels(surv_pre_post_ini[Pre_Temp!="cycling"])
surv_pre_post[,Wolb:=relevel(Wolb,"Wolb+")]

##Diagnostics
```

```r
#How many individuals were tested
ftable(xtabs(~Pre_Temp+Post_Temp+Wolb,surv_pre_post))
```

```
##                    Wolb Wolb+ Wolb-
## Pre_Temp Post_Temp
## 18       18              100   100
##          25              100   100
## 25       18              100   100
##          25              100   100
```

```r
ftable(xtabs(~Pre_Temp+Post_Temp+Wolb,surv_cyc))
```

```
##                    Wolb Wolb- Wolb+
## Pre_Temp Post_Temp
## cycling  cycling         100   100
```

```r
##Data analysis
# Full model
cox_pre_post_full<-coxme(Surv(Time,Status)~Wolb*Pre_Temp*Post_Temp+(1|RepFull),
                      surv_pre_post)
Anova(cox_pre_post_full)
```

```
## Analysis of Deviance Table (Type II tests)
##
## Response: Surv(Time, Status)
##                         Df   Chisq Pr(>Chisq)
## Wolb                     1 18.4874  1.710e-05 ***
## Pre_Temp                 1 16.3617  5.233e-05 ***
## Post_Temp                1 30.5079  3.325e-08 ***
## Wolb:Pre_Temp            1  7.0238   0.008043 **
## Wolb:Post_Temp           1  0.0049   0.944290
## Pre_Temp:Post_Temp       1  0.0596   0.807088
## Wolb:Pre_Temp:Post_Temp  1  2.3646   0.124117
## ---
## Signif. codes:  0 '***' 0.001 '**' 0.01 '*' 0.05 '.' 0.1 ' ' 1
```

```r
#simpler model
cox_pre_post_final<-coxme(Surv(Time,Status)~Wolb*Pre_Temp+Post_Temp+(1|RepFull),
                      surv_pre_post)
Anova(cox_pre_post_final)
```

```
## Analysis of Deviance Table (Type II tests)
##
## Response: Surv(Time, Status)
##               Df   Chisq Pr(>Chisq)
## Wolb           1 18.2611  1.926e-05 ***
## Pre_Temp       1 16.1596  5.822e-05 ***
## Post_Temp      1 29.8053  4.777e-08 ***
## Wolb:Pre_Temp  1  6.7722   0.009259 **
## ---
## Signif. codes:  0 '***' 0.001 '**' 0.01 '*' 0.05 '.' 0.1 ' ' 1
```

```r
summary(cox_pre_post_final)
```

```
## Cox mixed-effects model fit by maximum likelihood
##   Data: surv_pre_post
##   events, n = 500, 800
```

```
##   Iterations= 14 77
##                      NULL Integrated    Fitted
## Log-likelihood -3137.045  -2996.654 -2909.238
##
##                     Chisq    df p    AIC    BIC
## Integrated loglik 280.78  5.00 0 270.78 249.71
##  Penalized loglik 455.61 60.98 0 333.65  76.64
##
## Model:  Surv(Time, Status) ~ Wolb * Pre_Temp + Post_Temp + (1 | RepFull)
## Fixed coefficients
##                            coef exp(coef)  se(coef)     z       p
## WolbWolb-             0.3450515 1.4120627 0.2736339  1.26 2.1e-01
## Pre_Temp25           -1.3396287 0.2619429 0.2876196 -4.66 3.2e-06
## Post_Temp25           1.0147959 2.7588002 0.1858797  5.46 4.8e-08
## WolbWolb-:Pre_Temp25  1.0392922 2.8272153 0.3993688  2.60 9.3e-03
##
## Random effects
##  Group   Variable  Std Dev   Variance
##  RepFull Intercept 0.7535451 0.5678303
```

Significant interaction between *Wolbachia* and pre-infection temperature

```
#Hazards ratios between wolb+ and wolb- at both temp
pre_temp_Wolb<-lsmeans::lsmeans(cox_pre_post_final,pairwise~Wolb|Pre_Temp)
summary(pre_temp_Wolb,by=NULL,adj="holm")
```

```
## $lsmeans
##  Wolb  Pre_Temp lsmean    SE  df asymp.LCL asymp.UCL
##  Wolb+ 18        0.237 0.170 Inf    -0.187     0.661
##  Wolb- 18        0.583 0.168 Inf     0.162     1.003
##  Wolb+ 25       -1.102 0.179 Inf    -1.548    -0.656
##  Wolb- 25        0.282 0.170 Inf    -0.143     0.708
##
## Results are averaged over the levels of: Post_Temp
## Results are given on the log (not the response) scale.
## Confidence level used: 0.95
## Conf-level adjustment: bonferroni method for 4 estimates
##
## $contrasts
##  contrast      Pre_Temp estimate    SE  df z.ratio p.value
##  Wolb+ - Wolb- 18         -0.345 0.274 Inf -1.261  0.2073
##  Wolb+ - Wolb- 25         -1.384 0.287 Inf -4.823  <.0001
##
## Results are averaged over the levels of: Post_Temp
## Results are given on the log (not the response) scale.
## P value adjustment: holm method for 2 tests
```

*Wolbachia* only makes a difference if pre-infection temperature is 25ºC

```
#direct comparison of Wolb effect at different temp
contrast(pre_temp_Wolb$contrasts,"pairwise",by="contrast")
```

```
## contrast = Wolb+ - Wolb-:
##  contrast1 estimate    SE  df z.ratio p.value
##  18 - 25       1.04 0.399 Inf 2.602   0.0093
##
## Results are averaged over the levels of: Post_Temp
## Results are given on the log (not the response) scale.
```

*Wolbachia* effect is significantly different between pre-infection temperatures

```
#Hazards ratios between pre-temps when wolb is either present or not
Wolb_pre_temp<-lsmeans::lsmeans(cox_pre_post_final,pairwise~Pre_Temp|Wolb)
summary(Wolb_pre_temp,by=NULL,adj="holm")
```

```
## $lsmeans
##  Pre_Temp Wolb  lsmean    SE  df asymp.LCL asymp.UCL
##  18       Wolb+  0.237 0.170 Inf    -0.187     0.661
##  25       Wolb+ -1.102 0.179 Inf    -1.548    -0.656
##  18       Wolb-  0.583 0.168 Inf     0.162     1.003
##  25       Wolb-  0.282 0.170 Inf    -0.143     0.708
##
## Results are averaged over the levels of: Post_Temp
## Results are given on the log (not the response) scale.
## Confidence level used: 0.95
## Conf-level adjustment: bonferroni method for 4 estimates
##
## $contrasts
##  contrast Wolb  estimate    SE  df z.ratio p.value
##  18 - 25  Wolb+     1.34 0.288 Inf 4.658   <.0001
##  18 - 25  Wolb-     0.30 0.276 Inf 1.090   0.2759
##
## Results are averaged over the levels of: Post_Temp
## Results are given on the log (not the response) scale.
## P value adjustment: holm method for 2 tests
```

Pre-infection temperature only makes a difference if *Wolbachia* is present

##### Figures 2A and 2B

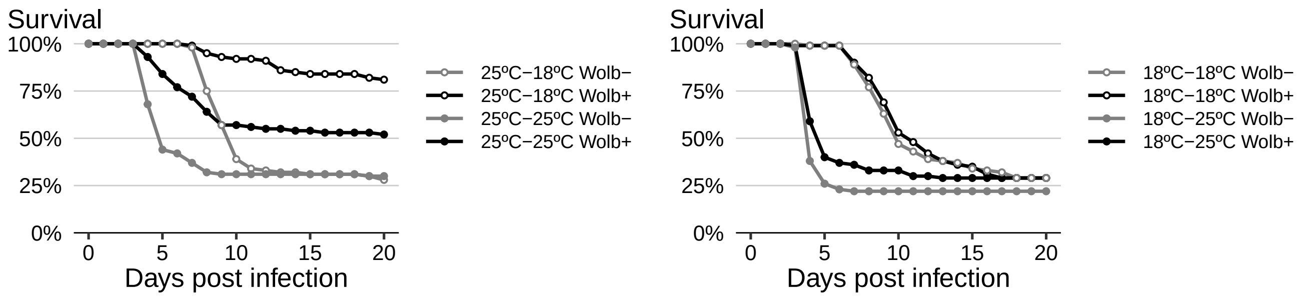

#### DCV levels and different pre-infection temperatures

```r
#load data
pre_DCV<-fread("dataset_s5.txt")
pre_DCV<-pre_DCV[,pre_wolb:=paste(Pre_inf,Wolb,sep="_")][,lapply(.SD,char_asfactor)]
pre_DCV<-pre_DCV[,logRatio:=log10(Ratio)]
pre_DCV$Pre_inf<-as.factor(as.character(pre_DCV$Pre_inf))

head(pre_DCV)
```

```
##    Pre_inf  Wolb Sample_name     Ratio Rep pre_wolb logRatio
## 1:      18 Wolb+    CS 18 10  662.4316   a 18_Wolb+ 2.821141
## 2:      18 Wolb+     CS 18 5 1045.3680   a 18_Wolb+ 3.019269
## 3:      18 Wolb+     CS 18 6  713.7923   a 18_Wolb+ 2.853572
## 4:      18 Wolb+     CS 18 7 1820.3450   a 18_Wolb+ 3.260154
## 5:      18 Wolb+     CS 18 8 1001.5730   a 18_Wolb+ 3.000683
## 6:      18 Wolb+     CS 18 9  539.2514   a 18_Wolb+ 2.731791
```

```r
#lmer

mod_DCV<-lmer(logRatio~Pre_inf*Wolb + (1|Rep),data=pre_DCV)
Anova(mod_DCV)
```

```
## Analysis of Deviance Table (Type II Wald chisquare tests)
##
## Response: logRatio
##                Chisq Df Pr(>Chisq)
## Pre_inf       45.093  1  1.879e-11 ***
## Wolb         145.361  1  < 2.2e-16 ***
## Pre_inf:Wolb  33.996  1  5.522e-09 ***
## ---
## Signif. codes:  0 '***' 0.001 '**' 0.01 '*' 0.05 '.' 0.1 ' ' 1
```

```r
summary(mod_DCV)
```

```
## Linear mixed model fit by REML. t-tests use Satterthwaite's method [
## lmerModLmerTest]
## Formula: logRatio ~ Pre_inf * Wolb + (1 | Rep)
##    Data: pre_DCV
##
## REML criterion at convergence: 148.4
##
## Scaled residuals:
##     Min      1Q  Median      3Q     Max
## -4.3523 -0.2458  0.1647  0.4542  2.2721
##
## Random effects:
##  Groups   Name        Variance Std.Dev.
##  Rep      (Intercept) 1.3033   1.142
##  Residual             0.5374   0.733
## Number of obs: 64, groups:  Rep, 2
##
## Fixed effects:
##                     Estimate Std. Error      df t value Pr(>|t|)
## (Intercept)           3.1264     0.8281  1.0776   3.775    0.151
## Pre_inf25            -0.1621     0.2592 59.0000  -0.625    0.534
```

```
## WolbWolb+            -1.1410     0.2592 59.0000  -4.402 4.56e-05 ***
## Pre_inf25:WolbWolb+  -2.1371     0.3665 59.0000  -5.831 2.47e-07 ***
## ---
## Signif. codes:  0 '***' 0.001 '**' 0.01 '*' 0.05 '.' 0.1 ' ' 1
##
## Correlation of Fixed Effects:
##             (Intr) Pr_n25 WlbWl+
## Pre_inf25   -0.156
## WolbWolb+   -0.156  0.500
## Pr_nf25:WW+  0.111 -0.707 -0.707
```

There is interaction between pre-infection temperature and *Wolbachia*

```
lsm_pre_DCV_wolb<-lsmeans::lsmeans(mod_DCV,pairwise~Wolb|Pre_inf,adj="none")
summary(lsm_pre_DCV_wolb,by=NULL,adj="holm")
```

```
## $lsmeans
##  Wolb  Pre_inf lsmean    SE   df lower.CL upper.CL
##  Wolb- 18       3.126 0.828 1.08    -29.1     35.4
##  Wolb+ 18       1.985 0.828 1.08    -30.3     34.3
##  Wolb- 25       2.964 0.828 1.08    -29.3     35.2
##  Wolb+ 25      -0.314 0.828 1.08    -32.6     32.0
##
## Degrees-of-freedom method: kenward-roger
## Confidence level used: 0.95
## Conf-level adjustment: bonferroni method for 4 estimates
##
## $contrasts
##  contrast      Pre_inf estimate    SE df t.ratio p.value
##  Wolb- - Wolb+ 18          1.14 0.259 59  4.402  <.0001
##  Wolb- - Wolb+ 25          3.28 0.259 59 12.648  <.0001
##
## Degrees-of-freedom method: kenward-roger
## P value adjustment: holm method for 2 tests
```

```
#fold changes
'^'(10,summary(lsm_pre_DCV_wolb,by=NULL,adj="holm")$contrasts$estimate)
```

```
## [1]   13.83501 1896.84848
```

```
#direct comparison of Wolb effect at different temp
contrast(lsm_pre_DCV_wolb$contrasts,"pairwise",by="contrast")
```

```
## contrast = Wolb- - Wolb+:
##  contrast1 estimate    SE df t.ratio p.value
##  18 - 25      -2.14 0.367 59 -5.831  <.0001
##
## Degrees-of-freedom method: kenward-roger
```

*Wolbachia* increases resistance at both pre-infection temperatures. But *Wolbachia* induces more resistance at pre-infection temperature of 25ºC

```r
#differences between pre-temp in absence or presence of Wolb
lsm_pre_DCV_temp<-lsmeans::lsmeans(mod_DCV,pairwise~Pre_inf|Wolb,adj="none")
summary(lsm_pre_DCV_temp,by=NULL,adj="holm")
```

```
## $lsmeans
##  Pre_inf Wolb  lsmean    SE   df lower.CL upper.CL
##  18      Wolb-  3.126 0.828 1.08    -29.1     35.4
##  25      Wolb-  2.964 0.828 1.08    -29.3     35.2
##  18      Wolb+  1.985 0.828 1.08    -30.3     34.3
##  25      Wolb+ -0.314 0.828 1.08    -32.6     32.0
##
## Degrees-of-freedom method: kenward-roger
## Confidence level used: 0.95
## Conf-level adjustment: bonferroni method for 4 estimates
##
## $contrasts
##  contrast Wolb  estimate    SE df t.ratio p.value
##  18 - 25  Wolb-    0.162 0.259 59 0.625   0.5341
##  18 - 25  Wolb+    2.299 0.259 59 8.871   <.0001
##
## Degrees-of-freedom method: kenward-roger
## P value adjustment: holm method for 2 tests
```

Pre-infection temperature only changes resistance when *Wolbachia* is present

##### Figures 2C and S1

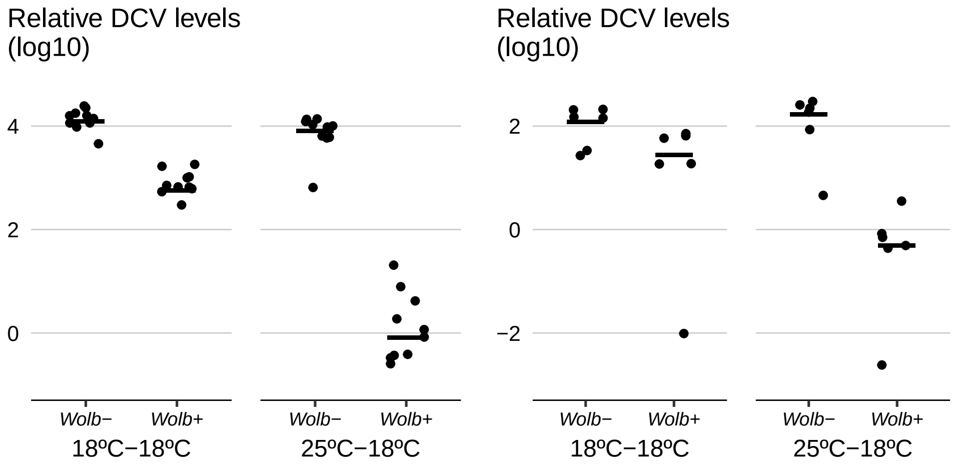

#### *Wolbachia* levels and different pre-infection temperatures

```r
##Fig 2D
#Wolbachia levels

pre_Wolb<-fread("dataset_s6.txt")[,lapply(.SD,char_asfactor)]
pre_Wolb[,Wolb:=rel_wolb][,logWolb:=log10(Wolb)]
```

```r
##Data Analysis
#Linear model

pre_wolb_lm<-lm(logWolb~Temp, data=pre_Wolb)
summary(pre_wolb_lm)
```

```
##
## Call:
## lm(formula = logWolb ~ Temp, data = pre_Wolb)
##
## Residuals:
##       Min        1Q    Median        3Q       Max
## -0.103011 -0.047381  0.002781  0.047650  0.094834
##
## Coefficients:
##              Estimate Std. Error t value Pr(>|t|)
## (Intercept) -0.002111   0.018891  -0.112    0.912
## Temp25C      0.155470   0.026716   5.819 1.63e-05 ***
## ---
## Signif. codes:  0 '***' 0.001 '**' 0.01 '*' 0.05 '.' 0.1 ' ' 1
##
## Residual standard error: 0.05974 on 18 degrees of freedom
## Multiple R-squared:  0.653,  Adjusted R-squared:  0.6337
## F-statistic: 33.87 on 1 and 18 DF,  p-value: 1.635e-05
```

```r
Anova(pre_wolb_lm)
```

```
## Anova Table (Type II tests)
##
## Response: logWolb
##             Sum Sq Df F value    Pr(>F)
## Temp      0.120854  1  33.866 1.635e-05 ***
## Residuals 0.064235 18
## ---
## Signif. codes:  0 '***' 0.001 '**' 0.01 '*' 0.05 '.' 0.1 ' ' 1
```

```r
#estimated fold differences in Wolbachia levels between pre-temp
'^'(10,pre_wolb_lm$coefficients)
```

```
## (Intercept)     Temp25C
##   0.9951519   1.4304396
```

```r
1 / '^'(10,pre_wolb_lm$coefficients)
```

```
## (Intercept)     Temp25C
##   1.0048717   0.6990858
```

*Wolbachia* levels differ between development temperatures, are higher at higher temperature

**Figures 2D**

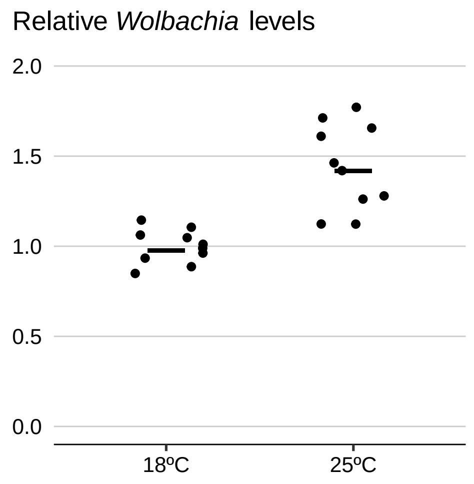

### Figure 3 and S2-4

#### Effect of pre-infection temperature on survival to DCV infection in different host and *Wolbachia* genetic backgrounds

```
#Figure 3 A and B
genotypes <- fread("dataset_s7.txt")

genotypes[,Gen:=gsub("c.*","",Gen)][,Gen:=tolower(Gen)][,RepFull:=paste(Gen,Dose,Preinf,Wolbachia,Test,R
genotypes[,PlotSeries:=paste(Gen,Test,sep="_")][,Condition:=paste(Wolbachia,Preinf)]
genotypes<-genotypes[,lapply(.SD,char_asfactor)][,Preinf:=as.factor(Preinf)]

genotypes_toplot<-data.table(fitsurv(.(Gen,Wolbachia,Preinf,Dose),genotypes))

##Diagnostics
#How many individuals were tested
ftable(xtabs(~Dose+Wolbachia+Gen+Test+Preinf,genotypes))
```

```
##                                 Preinf 18 25
## Dose Wolbachia Gen  Test
## E7   Wol-      alj1 01_12_2014         40 40
##                     27_11_2014         40 40
##                w20  01_12_2014         40 40
##                     27_11_2014         40 40
##      Wol+      alj1 01_12_2014         40 40
##                     27_11_2014         40 40
##                w20  01_12_2014         40 40
##                     27_11_2014         40 40
## E8   Wol-      alj1 01_12_2014         40 40
##                     27_11_2014         40 40
##                w20  01_12_2014         40 40
##                     27_11_2014         40 40
##      Wol+      alj1 01_12_2014         40 40
##                     27_11_2014         40 40
##                w20  01_12_2014         40 40
```

```
##                     27_11_2014        44 40
## E9   Wol-      alj1 01_12_2014        40 40
##                     27_11_2014        40 40
##                w20  01_12_2014        40 40
##                     27_11_2014        40 40
##      Wol+      alj1 01_12_2014        40 40
##                     27_11_2014        40 40
##                w20  01_12_2014        40 40
##                     27_11_2014        40 40
```

```
##Models

#Analyse one genotype at a time

#Aljezur flies, Cox model
#Full model
alj_all<-genotypes[Gen=="alj1"]

alj_cox_full<-coxme(Surv(Time,Status)~Wolbachia*Dose*Preinf+(1|Test/RepFull),data=alj_all)
Anova(alj_cox_full,test.statistic = "LR")
```

```
## Analysis of Deviance Table (Type II tests)
##                         LR Chisq Df Pr(>Chisq)
## Wolbachia                 22.498  1  2.104e-06 ***
## Dose                      50.914  2  8.793e-12 ***
## Preinf                     5.258  1    0.02184 *
## Wolbachia:Dose             3.009  2    0.22211
## Wolbachia:Preinf          27.037  1  1.996e-07 ***
## Dose:Preinf                0.729  2    0.69451
## Wolbachia:Dose:Preinf      1.407  2    0.49485
## ---
## Signif. codes:  0 '***' 0.001 '**' 0.01 '*' 0.05 '.' 0.1 ' ' 1
```

```
#simplified model
alj_cox_model2<-coxme(Surv(Time,Status)~Wolbachia*Preinf+Dose+(1|Test/RepFull),data=alj_all)
Anova(alj_cox_model2,test.statistic = "LR")
```

```
## Analysis of Deviance Table (Type II tests)
##                    LR Chisq Df Pr(>Chisq)
## Wolbachia            22.376  1  2.242e-06 ***
## Preinf                5.223  1     0.0223 *
## Dose                 50.914  2  8.793e-12 ***
## Wolbachia:Preinf     26.183  1  3.105e-07 ***
## ---
## Signif. codes:  0 '***' 0.001 '**' 0.01 '*' 0.05 '.' 0.1 ' ' 1
```

```
summary(alj_cox_model2)
```

```
## Cox mixed-effects model fit by maximum likelihood
##   Data: alj_all
##   events, n = 842, 960
##   Iterations= 20 105
##                     NULL Integrated    Fitted
## Log-likelihood -5188.363   -5049.27 -4982.198
##
##                       Chisq    df p    AIC    BIC
```

```
## Integrated loglik 278.19  7.00 0 264.19 231.03
##  Penalized loglik 412.33 55.92 0 300.48  35.64
##
## Model:  Surv(Time, Status) ~ Wolbachia * Preinf + Dose + (1 | Test/RepFull)
## Fixed coefficients
##                                coef exp(coef)  se(coef)     z       p
## WolbachiaWol+           -0.04456881 0.9564098 0.1475228 -0.30 7.6e-01
## Preinf25                 0.29156245 1.3385172 0.1483443  1.97 4.9e-02
## DoseE8                   0.57182227 1.7714923 0.1307041  4.37 1.2e-05
## DoseE9                   1.05424879 2.8698185 0.1299828  8.11 5.6e-16
## WolbachiaWol+:Preinf25 -1.14555241 0.3180482 0.2109993 -5.43 5.7e-08
##
## Random effects
##  Group        Variable    Std Dev    Variance
##  Test/RepFull (Intercept) 0.38367135 0.14720370
##  Test         (Intercept) 0.17308333 0.02995784
```

Interaction between *Wolbachia* and temperature in Aljezur1 flies

```
#Test the effect of wolbachia at different pre-infection temperature
alj_cox_contr=lsmeans::lsmeans(alj_cox_model2,pairwise~Wolbachia|Preinf)
summary(alj_cox_contr$contrasts,by=NULL,adj="holm")
```

```
##  contrast    Preinf estimate    SE  df z.ratio p.value
##  Wol- - Wol+ 18       0.0446 0.148 Inf 0.302   0.7626
##  Wol- - Wol+ 25       1.1901 0.151 Inf 7.898   <.0001
##
## Results are averaged over the levels of: Dose
## Results are given on the log (not the response) scale.
## P value adjustment: holm method for 2 tests
```

```
#direct comparison of Wolb effect at different temp
contrast(alj_cox_contr$contrasts,"pairwise",by="contrast")
```

```
## contrast = Wol- - Wol+:
##  contrast1 estimate    SE  df z.ratio p.value
##  18 - 25      -1.15 0.211 Inf -5.429  <.0001
##
## Results are averaged over the levels of: Dose
## Results are given on the log (not the response) scale.
```

*Wolbachia* only has an effect if flies raised at 25ºC

```
#W20  flies, Cox model
#Full model
w20_all<-genotypes[Gen=="w20"]

w20_cox_full<-coxme(Surv(Time,Status)~Wolbachia*Dose*Preinf+(1|Test/RepFull),data=w20_all)
Anova(w20_cox_full,test.statistic = "LR")
```

```
## Analysis of Deviance Table (Type II tests)
##                        LR Chisq Df Pr(>Chisq)
```

```
## Wolbachia               80.541  1  < 2.2e-16 ***
## Dose                    53.989  2  1.889e-12 ***
## Preinf                  19.556  1  9.771e-06 ***
## Wolbachia:Dose           1.083  2     0.5819
## Wolbachia:Preinf        29.042  1  7.081e-08 ***
## Dose:Preinf              0.777  2     0.6779
## Wolbachia:Dose:Preinf    0.836  2     0.6584
## ---
## Signif. codes:  0 '***' 0.001 '**' 0.01 '*' 0.05 '.' 0.1 ' ' 1
```

```
#Simplified model
w20_cox_model2<-coxme(Surv(Time,Status)~Wolbachia*Preinf+Dose+(1|Test/RepFull),data=w20_all)
Anova(w20_cox_model2,test.statistic = "LR")
```

```
## Analysis of Deviance Table (Type II tests)
##                  LR Chisq Df Pr(>Chisq)
## Wolbachia          80.263  1  < 2.2e-16 ***
## Preinf             19.451  1  1.032e-05 ***
## Dose               53.989  2  1.889e-12 ***
## Wolbachia:Preinf   28.363  1  1.006e-07 ***
## ---
## Signif. codes:  0 '***' 0.001 '**' 0.01 '*' 0.05 '.' 0.1 ' ' 1
```

```
summary(w20_cox_model2)
```

```
## Cox mixed-effects model fit by maximum likelihood
##   Data: w20_all
##   events, n = 734, 964
##   Iterations= 7 51
##                     NULL Integrated    Fitted
## Log-likelihood -4639.69  -4246.389 -4142.787
##
##                     Chisq    df p    AIC    BIC
## Integrated loglik 786.60  7.00 0 772.60 740.41
##  Penalized loglik 993.81 72.38 0 849.04 516.18
##
## Model:  Surv(Time, Status) ~ Wolbachia * Preinf + Dose + (1 | Test/RepFull)
## Fixed coefficients
##                             coef exp(coef)  se(coef)     z       p
## WolbachiaWol+          -1.21292804 0.2973254 0.2127377 -5.70 1.2e-08
## Preinf25                0.01231417 1.0123903 0.2240180  0.05 9.6e-01
## DoseE8                  1.18444422 3.2688696 0.2018180  5.87 4.4e-09
## DoseE9                  1.60143882 4.9601641 0.1982039  8.08 6.7e-16
## WolbachiaWol+:Preinf25 -1.80380488 0.1646711 0.3303489 -5.46 4.8e-08
##
## Random effects
##  Group        Variable    Std Dev      Variance
##  Test/RepFull (Intercept) 0.6762892716 0.4573671789
##  Test         (Intercept) 0.0201313478 0.0004052712
```

Interaction between *Wolbachia* and temperature in W20 flies

```
#Test the effect of wolbachia at different pre-infection temperature (minimal model)**
w20_cox_contr=lsmeans::lsmeans(w20_cox_model2,pairwise~Wolbachia|Preinf)
```

```r
summary(w20_cox_contr$contrasts,by=NULL,adj="holm")
```

```
##  contrast    Preinf estimate    SE  df z.ratio p.value
##  Wol- - Wol+ 18         1.21 0.213 Inf  5.702  <.0001
##  Wol- - Wol+ 25         3.02 0.235 Inf 12.821  <.0001
##
## Results are averaged over the levels of: Dose
## Results are given on the log (not the response) scale.
## P value adjustment: holm method for 2 tests
```

```r
#direct comparison of Wolb effect at different temp
contrast(w20_cox_contr$contrasts,"pairwise",by="contrast")
```

```
## contrast = Wol- - Wol+:
##  contrast1 estimate   SE  df z.ratio p.value
##  18 - 25       -1.8 0.33 Inf -5.460  <.0001
##
## Results are averaged over the levels of: Dose
## Results are given on the log (not the response) scale.
```

*Wolbachia* has an effect at both temperatures, effect is stronger in flies raised at 25ºC

##### Figures 3A, 3B, S2A, S2B

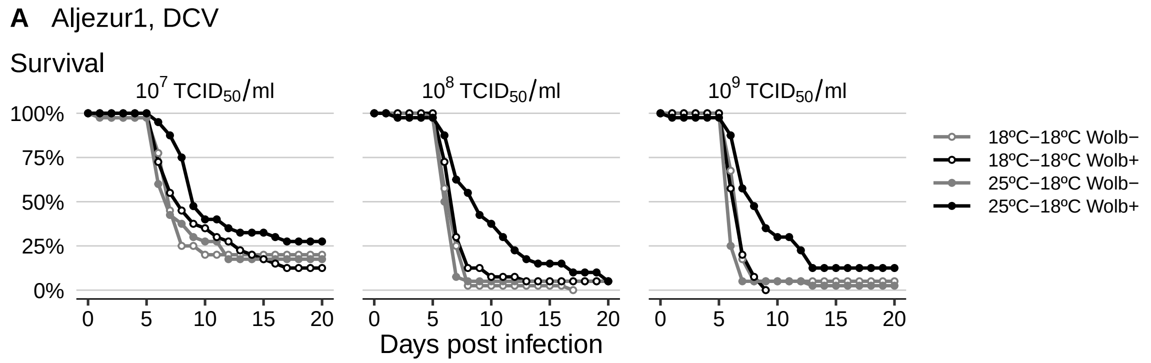

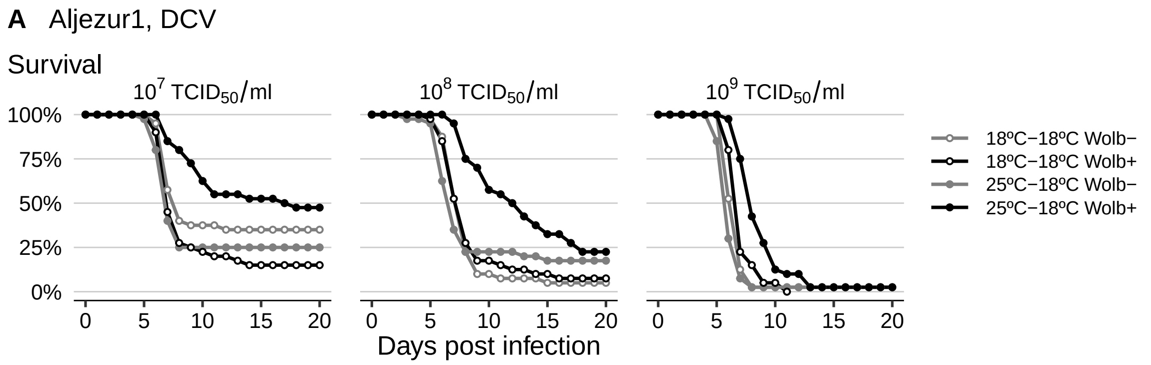

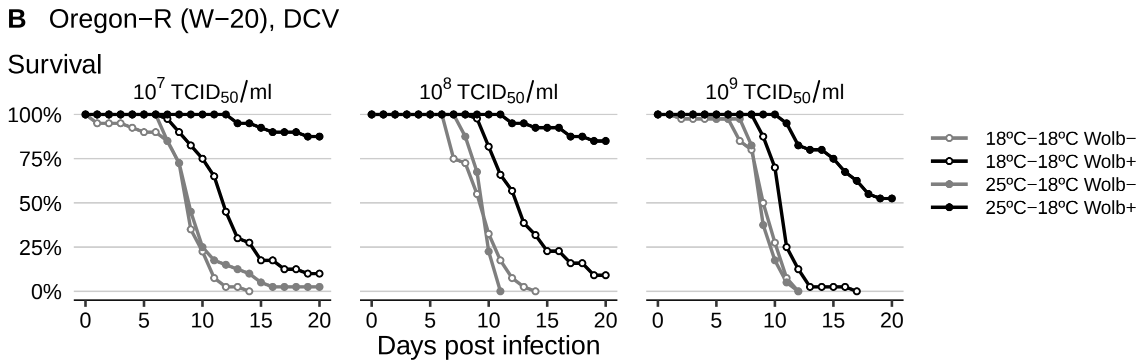

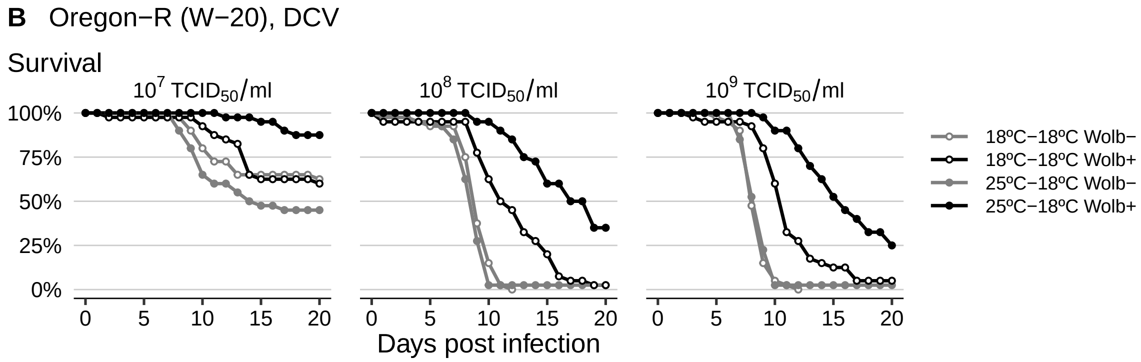

#### Effect of pre-infection temperature on survival to FHV infection

```
###Effect of Pre-temp on FHV
##Data import
surv_FHV<-fread("dataset_s8.txt")

surv_FHV<-surv_FHV[,Preinf:=as.factor(Preinf)]
surv_FHV[,RepFull:=interaction(Preinf,Wolbachia,Dose,Replicate,sep = "_")][,Condition:=paste(Wolbachia,
surv_FHV<-surv_FHV[,lapply(.SD,char_asfactor)]

##Diagnostics
#How many individuals were tested
ftable(xtabs(~Test+Preinf+Dose+Wolbachia,surv_FHV))
```

```
##                         Wolbachia Wol- Wol+
## Test       Preinf Dose
## 09_02_2015 18     E6                50   51
##                   E7                50   50
##                   E8                49   50
##                   E9                 0    0
##            25     E6                49   48
##                   E7                50   50
##                   E8                50   50
##                   E9                 0    0
## 21_11_2014 18     E6                 0    0
##                   E7                50   50
##                   E8                50   50
##                   E9                50   50
##            25     E6                 0    0
```

```
##                    E7                50   50
##                    E8                50   50
##                    E9                50   50
```

```
#Kill one fly of wolb+ 25-18 E6 sample, in the last day, to allow model to converge
surv_FHVa <- surv_FHV
surv_FHVa$Status[which(surv_FHVa$Genotype == "CS 25-18 E6")[1]]<-1

#Data analysis
#Cox models
#Full model
surv_FHV_cox_full<-coxme(Surv(Time,Status)~(Wolbachia*Dose*Preinf)+(1|Test/RepFull),
                    data=surv_FHVa)
Anova(surv_FHV_cox_full,test.statistic = "LR")
```

```
## Analysis of Deviance Table (Type II tests)
##                        LR Chisq Df Pr(>Chisq)
## Wolbachia               102.003  1  < 2.2e-16 ***
## Dose                    194.917  3  < 2.2e-16 ***
## Preinf                   48.565  1  3.195e-12 ***
## Wolbachia:Dose            4.409  3    0.22056
## Wolbachia:Preinf        143.421  1  < 2.2e-16 ***
## Dose:Preinf               0.958  3    0.81143
## Wolbachia:Dose:Preinf    10.825  3    0.01271 *
## ---
## Signif. codes:  0 '***' 0.001 '**' 0.01 '*' 0.05 '.' 0.1 ' ' 1
```

There is a significant interaction between pre-infection temperature, *Wolbachia*, and dose of infection

```
#effect of Wolb at different temp and doses
surv_FHV_temp_contr=lsmeans::lsmeans(surv_FHV_cox_full,pairwise~Wolbachia|Preinf:Dose)
summary(surv_FHV_temp_contr,by=NULL,adj="holm")
```

```
## $lsmeans
##  Wolbachia Preinf Dose lsmean    SE  df asymp.LCL asymp.UCL
##  Wol-      18     E6   -1.772 0.313 Inf    -2.698    -0.846
##  Wol+      18     E6   -3.193 0.565 Inf    -4.864    -1.522
##  Wol-      25     E6   -1.847 0.326 Inf    -2.812    -0.882
##  Wol+      25     E6   -4.268 0.967 Inf    -7.125    -1.411
##  Wol-      18     E7    1.195 0.139 Inf     0.785     1.604
##  Wol+      18     E7    0.188 0.153 Inf    -0.264     0.641
##  Wol-      25     E7    1.134 0.138 Inf     0.725     1.542
##  Wol+      25     E7   -2.717 0.387 Inf    -3.862    -1.572
##  Wol-      18     E8    2.370 0.140 Inf     1.957     2.782
##  Wol+      18     E8    1.598 0.136 Inf     1.197     1.999
##  Wol-      25     E8    2.406 0.140 Inf     1.992     2.820
##  Wol+      25     E8   -3.132 0.469 Inf    -4.517    -1.747
##  Wol-      18     E9    2.441 0.188 Inf     1.887     2.995
##  Wol+      18     E9    2.032 0.187 Inf     1.479     2.585
##  Wol-      25     E9    2.210 0.187 Inf     1.657     2.762
##  Wol+      25     E9   -1.780 0.567 Inf    -3.455    -0.106
##
## Results are given on the log (not the response) scale.
```

```
## Confidence level used: 0.95
## Conf-level adjustment: bonferroni method for 16 estimates
##
## $contrasts
##  contrast    Preinf Dose estimate    SE  df z.ratio p.value
##  Wol- - Wol+ 18     E6      1.420 0.660 Inf   2.153  0.0648
##  Wol- - Wol+ 25     E6      2.421 1.054 Inf   2.297  0.0648
##  Wol- - Wol+ 18     E7      1.006 0.186 Inf   5.405  <.0001
##  Wol- - Wol+ 25     E7      3.851 0.432 Inf   8.920  <.0001
##  Wol- - Wol+ 18     E8      0.772 0.167 Inf   4.632  <.0001
##  Wol- - Wol+ 25     E8      5.538 0.523 Inf  10.596  <.0001
##  Wol- - Wol+ 18     E9      0.409 0.229 Inf   1.786  0.0742
##  Wol- - Wol+ 25     E9      3.990 0.606 Inf   6.581  <.0001
##
## Results are given on the log (not the response) scale.
## P value adjustment: holm method for 8 tests
```

```
#interaction of Wolb and temp at different doses
surv_FHV_temp_contr_inter=contrast(surv_FHV_temp_contr$contrasts,by=c("contrast","Dose"),method="pairwi
summary(surv_FHV_temp_contr_inter,by=NULL,adj="holm")
```

```
##  contrast1 contrast    Dose estimate    SE  df z.ratio p.value
##  18 - 25   Wol- - Wol+ E6      -1.00 1.244 Inf -0.805  0.4208
##  18 - 25   Wol- - Wol+ E7      -2.84 0.469 Inf -6.069  <.0001
##  18 - 25   Wol- - Wol+ E8      -4.77 0.544 Inf -8.757  <.0001
##  18 - 25   Wol- - Wol+ E9      -3.58 0.647 Inf -5.536  <.0001
##
## Results are given on the log (not the response) scale.
## P value adjustment: holm method for 4 tests
```

*Wolbachia* effect is stronger at pre-infection temperature of 25ºC at all doses except $10^6$ $TCID_{50}$/ml

```
#effect of pre-infection temp with and without Wolb
summary(lsmeans::lsmeans(surv_FHV_cox_full,pairwise~Preinf|Dose:Wolbachia),by=NULL,adj = "holm")
```

```
## $lsmeans
##  Preinf Dose Wolbachia lsmean    SE  df asymp.LCL asymp.UCL
##  18     E6   Wol-      -1.772 0.313 Inf    -2.698    -0.846
##  25     E6   Wol-      -1.847 0.326 Inf    -2.812    -0.882
##  18     E7   Wol-       1.195 0.139 Inf     0.785     1.604
##  25     E7   Wol-       1.134 0.138 Inf     0.725     1.542
##  18     E8   Wol-       2.370 0.140 Inf     1.957     2.782
##  25     E8   Wol-       2.406 0.140 Inf     1.992     2.820
##  18     E9   Wol-       2.441 0.188 Inf     1.887     2.995
##  25     E9   Wol-       2.210 0.187 Inf     1.657     2.762
##  18     E6   Wol+      -3.193 0.565 Inf    -4.864    -1.522
##  25     E6   Wol+      -4.268 0.967 Inf    -7.125    -1.411
##  18     E7   Wol+       0.188 0.153 Inf    -0.264     0.641
##  25     E7   Wol+      -2.717 0.387 Inf    -3.862    -1.572
##  18     E8   Wol+       1.598 0.136 Inf     1.197     1.999
##  25     E8   Wol+      -3.132 0.469 Inf    -4.517    -1.747
##  18     E9   Wol+       2.032 0.187 Inf     1.479     2.585
##  25     E9   Wol+      -1.780 0.567 Inf    -3.455    -0.106
##
```

```
## Results are given on the log (not the response) scale.
## Confidence level used: 0.95
## Conf-level adjustment: bonferroni method for 16 estimates
##
## $contrasts
##  contrast Dose Wolbachia estimate    SE  df z.ratio p.value
##  18 - 25  E6   Wol-        0.0746 0.449 Inf  0.166  1.0000
##  18 - 25  E7   Wol-        0.0613 0.172 Inf  0.357  1.0000
##  18 - 25  E8   Wol-       -0.0364 0.162 Inf -0.224  1.0000
##  18 - 25  E9   Wol-        0.2314 0.227 Inf  1.021  1.0000
##  18 - 25  E6   Wol+        1.0756 1.159 Inf  0.928  1.0000
##  18 - 25  E7   Wol+        2.9055 0.436 Inf  6.660  <.0001
##  18 - 25  E8   Wol+        4.7299 0.519 Inf  9.109  <.0001
##  18 - 25  E9   Wol+        3.8121 0.606 Inf  6.292  <.0001
##
## Results are given on the log (not the response) scale.
## P value adjustment: holm method for 8 tests
```

Pre-infection temperature is not significant in the absence of *Wolbachia*. It is significant in the presence of *Wolbachia* at all doses except $10^6$ $\mathrm{TCID}_{50}$/ml.

**Figures 3C, S2C**

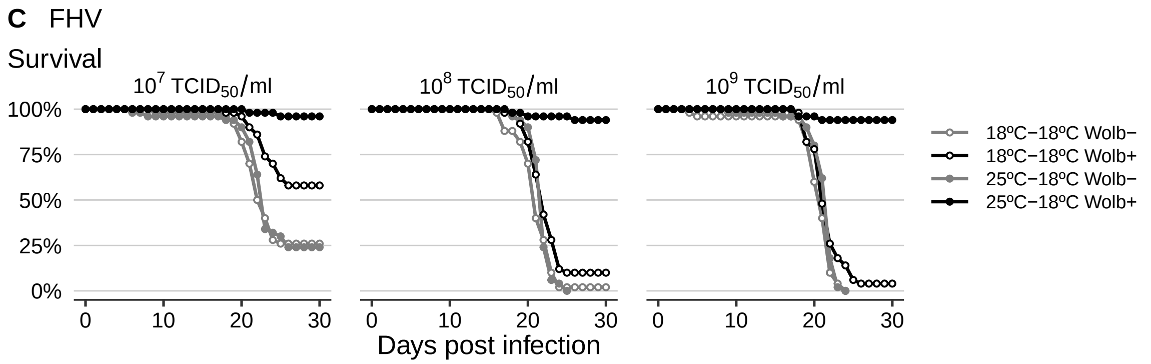

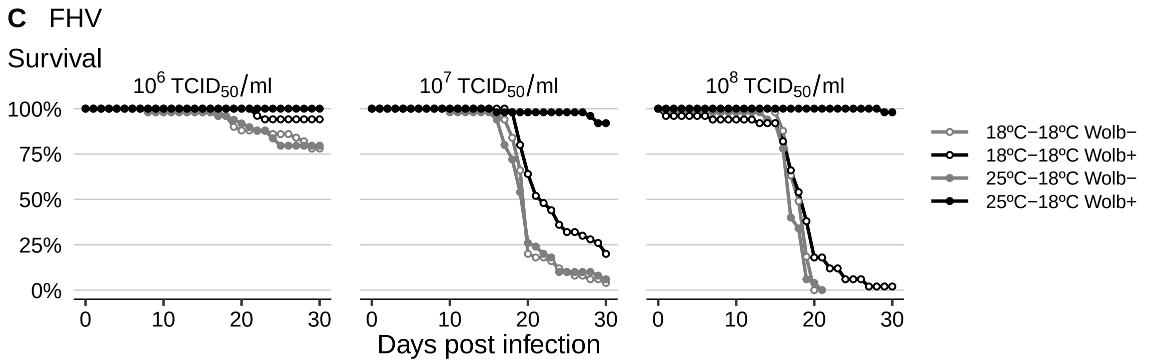

#### Survival with temperature cycling

```r
#Cox model
cox_cycling<-coxme(Surv(Time,Status)~Wolb+(1|RepFull),surv_cyc)

Anova(cox_cycling, test.statistic = "LR")
```

```
## Analysis of Deviance Table
##  Cox model: response is Surv(Time, Status)
## Terms added sequentially (first to last)
##
##       loglik  Chisq Df Pr(>|Chi|)
## NULL -737.91
## Wolb -725.48 24.867  1  6.142e-07 ***
## ---
## Signif. codes:  0 '***' 0.001 '**' 0.01 '*' 0.05 '.' 0.1 ' ' 1
```

```r
summary(cox_cycling)
```

```
## Cox mixed-effects model fit by maximum likelihood
##   Data: surv_cyc
##   events, n = 156, 200
##   Iterations= 7 32
##                     NULL Integrated    Fitted
## Log-likelihood -737.9147  -725.4812 -709.5113
##
##                   Chisq    df           p   AIC   BIC
## Integrated loglik 24.87  2.00 3.9828e-06 20.87 14.77
##  Penalized loglik 56.81 12.61 1.4132e-07 31.60 -6.85
##
## Model:  Surv(Time, Status) ~ Wolb + (1 | RepFull)
## Fixed coefficients
##                  coef exp(coef)  se(coef)     z     p
## WolbWolb+ -0.6671881 0.5131495 0.2719185 -2.45 0.014
##
## Random effects
##  Group   Variable  Std Dev   Variance
##  RepFull Intercept 0.4861021 0.2362953
```

*Wolbachia* increases survival in cycling temperatures

##### Figure S3

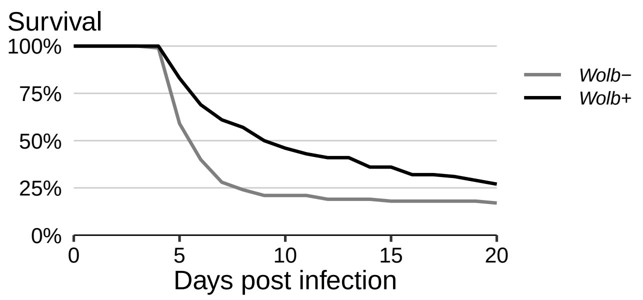

##### Effect of development and aging temperature before infection on survival

```
Surv_aging <- fread("dataset_s9.txt")[,lapply(.SD,char_asfactor)]
Surv_aging<-Surv_aging[,Rep:=paste(Test,Wolbachia, Development_temperature, Aging_temperature,Replicate
Surv_aging$Development_temperature<-as.factor(as.character(Surv_aging$Development_temperature))
Surv_aging$Aging_temperature<-as.factor(as.character(Surv_aging$Aging_temperature))

#Tested individuals
xtabs(~Wolbachia+RepTemp+Test,Surv_aging)
```

```
## , , Test = A
##
##          RepTemp
## Wolbachia 18_18 18_25 25_18 25_25
##      Wol-    50    50    50    50
##      Wol+    50    50    50    49
##
## , , Test = B
##
##          RepTemp
## Wolbachia 18_18 18_25 25_18 25_25
##      Wol-    50    50    50    50
##      Wol+    50    50    50    50
##
## , , Test = C
##
##          RepTemp
## Wolbachia 18_18 18_25 25_18 25_25
##      Wol-    50    50    50    50
##      Wol+    50    50    50    50
```

```
#full model
cox_Surv_aging_full<-coxme(Surv(Time,Status)~Wolbachia*Development_temperature*Aging_temperature+(1|Tes

Anova(cox_Surv_aging_full,test.statistic = "LR")
```

```
## Analysis of Deviance Table (Type II tests)
##                                                   LR Chisq Df Pr(>Chisq)
## Wolbachia                                           75.774  1  < 2.2e-16 ***
## Development_temperature                             29.275  1  6.279e-08 ***
## Aging_temperature                                    0.035  1    0.85061
## Wolbachia:Development_temperature                   40.163  1  2.336e-10 ***
## Wolbachia:Aging_temperature                          6.521  1    0.01066 *
## Development_temperature:Aging_temperature            0.633  1    0.42637
## Wolbachia:Development_temperature:Aging_temperature  0.513  1    0.47375
## ---
## Signif. codes:  0 '***' 0.001 '**' 0.01 '*' 0.05 '.' 0.1 ' ' 1
```

```
#simpler model
cox_Surv_aging_simple <-coxme(Surv(Time,Status)~Wolbachia*Aging_temperature+Wolbachia*Development_tempe
Anova(cox_Surv_aging_simple,test.statistic = "LR")
```

```
## Analysis of Deviance Table (Type II tests)
##                                   LR Chisq Df Pr(>Chisq)
## Wolbachia                           75.576  1  < 2.2e-16 ***
```

```
## Aging_temperature                        0.035  1    0.85061
## Development_temperature                 29.275  1  6.279e-08 ***
## Wolbachia:Aging_temperature              6.348  1    0.01175 *
## Wolbachia:Development_temperature       39.902  1  2.671e-10 ***
## ---
## Signif. codes:  0 '***' 0.001 '**' 0.01 '*' 0.05 '.' 0.1 ' ' 1
```

```
summary(cox_Surv_aging_simple)
```

```
## Cox mixed-effects model fit by maximum likelihood
##   Data: Surv_aging
##   events, n = 1064, 1199
##   Iterations= 12 64
##                      NULL Integrated    Fitted
## Log-likelihood -6774.882  -6532.167 -6463.197
##
##                     Chisq    df p    AIC    BIC
## Integrated loglik 485.43  7.00 0 471.43 436.64
##  Penalized loglik 623.37 58.64 0 506.08 214.63
##
## Model:  Surv(Time, Status) ~ Wolbachia * Aging_temperature + Wolbachia *      Development_temperatur
## Fixed coefficients
##                                                  coef exp(coef)  se(coef)     z
## WolbachiaWol+                             -0.237051814 0.7889504 0.1447667 -1.64
## Aging_temperature25                        0.222724245 1.2494760 0.1175657  1.89
## Development_temperature25                 -0.001842135 0.9981596 0.1174962 -0.02
## WolbachiaWol+:Aging_temperature25         -0.431474262 0.6495508 0.1699900 -2.54
## WolbachiaWol+:Development_temperature25 -1.158294456 0.3140213 0.1713643 -6.76
##                                                 p
## WolbachiaWol+                             1.0e-01
## Aging_temperature25                       5.8e-02
## Development_temperature25                 9.9e-01
## WolbachiaWol+:Aging_temperature25         1.1e-02
## WolbachiaWol+:Development_temperature25 1.4e-11
##
## Random effects
##  Group    Variable    Std Dev    Variance
##  Test/Rep (Intercept) 0.31406601 0.09863746
##  Test     (Intercept) 0.48075177 0.23112226
```

There is a significant interaction between *Wolbachia* and aging temperature, and between *Wolbachia* and development temperature.

```
#Contrasts

#Contrast between with and without Wolbachia at different development temperatures
contr_Wolb_Dev<-lsmeans::lsmeans(cox_Surv_aging_simple,pairwise~Wolbachia|Development_temperature)
summary(contr_Wolb_Dev)
```

```
## $lsmeans
## Development_temperature = 18:
##  Wolbachia  lsmean     SE  df asymp.LCL asymp.UCL
##  Wol-       0.5159 0.0728 Inf    0.3732     0.659
```

```
##  Wol+       0.0631 0.0727 Inf   -0.0795     0.206
##
## Development_temperature = 25:
##  Wolbachia  lsmean     SE  df asymp.LCL asymp.UCL
##  Wol-       0.5140 0.0730 Inf    0.3710     0.657
##  Wol+      -1.0970 0.0785 Inf   -1.2510    -0.943
##
## Results are averaged over the levels of: Aging_temperature
## Results are given on the log (not the response) scale.
## Confidence level used: 0.95
##
## $contrasts
## Development_temperature = 18:
##  contrast    estimate    SE  df z.ratio p.value
##  Wol- - Wol+    0.453 0.118 Inf  3.838  0.0001
##
## Development_temperature = 25:
##  contrast    estimate    SE  df z.ratio p.value
##  Wol- - Wol+    1.611 0.125 Inf 12.858  <.0001
##
## Results are averaged over the levels of: Aging_temperature
## Results are given on the log (not the response) scale.
```

```
contrast(contr_Wolb_Dev$contrasts,"pairwise",by="contrast")
```

```
## contrast = Wol- - Wol+:
##  contrast1 estimate    SE  df z.ratio p.value
##  18 - 25      -1.16 0.171 Inf -6.759  <.0001
##
## Results are averaged over the levels of: Aging_temperature
## Results are given on the log (not the response) scale.
```

*Wolbachia* has an effect at both development temperatures but effect is higher at 25ºC

```
#Contrast between with and without Wolbachia at different aging temperatures
contr_Wolb_Aging<-lsmeans::lsmeans(cox_Surv_aging_simple,pairwise~Wolbachia|Aging_temperature)
summary(contr_Wolb_Aging)
```

```
## $lsmeans
## Aging_temperature = 18:
##  Wolbachia lsmean     SE  df asymp.LCL asymp.UCL
##  Wol-       0.404 0.0728 Inf     0.261     0.546
##  Wol+      -0.413 0.0741 Inf    -0.558    -0.267
##
## Aging_temperature = 25:
##  Wolbachia lsmean     SE  df asymp.LCL asymp.UCL
##  Wol-       0.626 0.0731 Inf     0.483     0.770
##  Wol+      -0.621 0.0760 Inf    -0.770    -0.472
##
## Results are averaged over the levels of: Development_temperature
## Results are given on the log (not the response) scale.
## Confidence level used: 0.95
##
```

```
## $contrasts
## Aging_temperature = 18:
##  contrast    estimate    SE  df z.ratio p.value
##  Wol- - Wol+    0.816 0.120 Inf  6.806  <.0001
##
## Aging_temperature = 25:
##  contrast    estimate    SE  df z.ratio p.value
##  Wol- - Wol+    1.248 0.122 Inf 10.186  <.0001
##
## Results are averaged over the levels of: Development_temperature
## Results are given on the log (not the response) scale.
```

```
contrast(contr_Wolb_Aging$contrasts,"pairwise",by="contrast")
```

```
## contrast = Wol- - Wol+:
##  contrast1 estimate   SE  df z.ratio p.value
##  18 - 25     -0.431 0.17 Inf -2.538  0.0111
##
## Results are averaged over the levels of: Development_temperature
## Results are given on the log (not the response) scale.
```

*Wolbachia* has an effect at both aging temperatures but effect is higher at 25ºC

```
#Contrast between with and without Wolbachia at different development temperatures and aging temperatur
contr_Wolb_dev_temp_Aging<-lsmeans::lsmeans(cox_Surv_aging_simple,pairwise~Wolbachia|Development_tempera
summary(contr_Wolb_dev_temp_Aging,adj="holm",by=NULL)
```

```
## $lsmeans
##  Wolbachia Development_temperature Aging_temperature  lsmean     SE  df
##  Wol-      18                      18                 0.4045 0.0931 Inf
##  Wol+      18                      18                 0.1675 0.0951 Inf
##  Wol-      25                      18                 0.4027 0.0940 Inf
##  Wol+      25                      18                -0.9927 0.0982 Inf
##  Wol-      18                      25                 0.6273 0.0941 Inf
##  Wol+      18                      25                -0.0413 0.0951 Inf
##  Wol-      25                      25                 0.6254 0.0934 Inf
##  Wol+      25                      25                -1.2014 0.1010 Inf
##  asymp.LCL asymp.UCL
##     0.1500     0.659
##    -0.0927     0.428
##     0.1457     0.660
##    -1.2612    -0.724
##     0.3700     0.884
##    -0.3013     0.219
##     0.3699     0.881
##    -1.4777    -0.925
##
## Results are given on the log (not the response) scale.
## Confidence level used: 0.95
## Conf-level adjustment: bonferroni method for 8 estimates
##
## $contrasts
##  contrast    Development_temperature Aging_temperature estimate    SE  df
```

```
##   Wol- - Wol+ 18                      18                    0.237 0.145 Inf
##   Wol- - Wol+ 25                      18                    1.395 0.150 Inf
##   Wol- - Wol+ 18                      25                    0.669 0.146 Inf
##   Wol- - Wol+ 25                      25                    1.827 0.153 Inf
##   z.ratio p.value
##    1.637  0.1015
##    9.305  <.0001
##    4.578  <.0001
##   11.953  <.0001
##
## Results are given on the log (not the response) scale.
## P value adjustment: holm method for 4 tests
```

```
contrast(contr_Wolb_dev_temp_Aging$contrasts,"pairwise",by="contrast")
```

```
## contrast = Wol- - Wol+:
##   contrast1     estimate    SE  df z.ratio p.value
##   18,18 - 25,18   -1.158 0.171 Inf -6.759  <.0001
##   18,18 - 18,25   -0.431 0.170 Inf -2.538  0.0542
##   18,18 - 25,25   -1.590 0.242 Inf -6.558  <.0001
##   25,18 - 18,25    0.727 0.240 Inf  3.024  0.0133
##   25,18 - 25,25   -0.431 0.170 Inf -2.538  0.0542
##   18,25 - 25,25   -1.158 0.171 Inf -6.759  <.0001
##
## Results are given on the log (not the response) scale.
## P value adjustment: tukey method for comparing a family of 4 estimates
```

```
#Contrast between development temperatures with and without Wolbachia
contr_Dev_Wolb<-lsmeans::lsmeans(cox_Surv_aging_simple,pairwise~Development_temperature|Wolbachia)
summary(contr_Dev_Wolb)
```

```
## $lsmeans
## Wolbachia = Wol-:
##   Development_temperature  lsmean     SE  df asymp.LCL asymp.UCL
##   18                       0.5159 0.0728 Inf    0.3732     0.659
##   25                       0.5140 0.0730 Inf    0.3710     0.657
##
## Wolbachia = Wol+:
##   Development_temperature  lsmean     SE  df asymp.LCL asymp.UCL
##   18                       0.0631 0.0727 Inf   -0.0795     0.206
##   25                      -1.0970 0.0785 Inf   -1.2510    -0.943
##
## Results are averaged over the levels of: Aging_temperature
## Results are given on the log (not the response) scale.
## Confidence level used: 0.95
##
## $contrasts
## Wolbachia = Wol-:
##   contrast estimate    SE  df z.ratio p.value
##   18 - 25   0.00184 0.117 Inf 0.016   0.9875
##
## Wolbachia = Wol+:
##   contrast estimate    SE  df z.ratio p.value
##   18 - 25   1.16014 0.124 Inf 9.338   <.0001
##
```

```
## Results are averaged over the levels of: Aging_temperature
## Results are given on the log (not the response) scale.
```

Development temperature only has an effect in the presence of *Wolbachia*

```r
#Contrast between aging temperatures with and without Wolbachia
contr_Aging_Wolb <-lsmeans::lsmeans(cox_Surv_aging_simple,pairwise~Aging_temperature|Wolbachia)
summary(contr_Aging_Wolb)
```

```
## $lsmeans
## Wolbachia = Wol-:
##  Aging_temperature lsmean     SE  df asymp.LCL asymp.UCL
##  18                 0.404 0.0728 Inf     0.261     0.546
##  25                 0.626 0.0731 Inf     0.483     0.770
##
## Wolbachia = Wol+:
##  Aging_temperature lsmean     SE  df asymp.LCL asymp.UCL
##  18                -0.413 0.0741 Inf    -0.558    -0.267
##  25                -0.621 0.0760 Inf    -0.770    -0.472
##
## Results are averaged over the levels of: Development_temperature
## Results are given on the log (not the response) scale.
## Confidence level used: 0.95
##
## $contrasts
## Wolbachia = Wol-:
##  contrast estimate    SE  df z.ratio p.value
##  18 - 25    -0.223 0.118 Inf -1.894  0.0582
##
## Wolbachia = Wol+:
##  contrast estimate    SE  df z.ratio p.value
##  18 - 25     0.209 0.123 Inf  1.703  0.0886
##
## Results are averaged over the levels of: Development_temperature
## Results are given on the log (not the response) scale.
```

Aging temperature does not have an effect in direct contrasts. The interaction significance comes from slightly deleterious effect of higher temperature in the absence of *Wolbachia* and slightly beneficial effect in its presence.

##### Figure S4

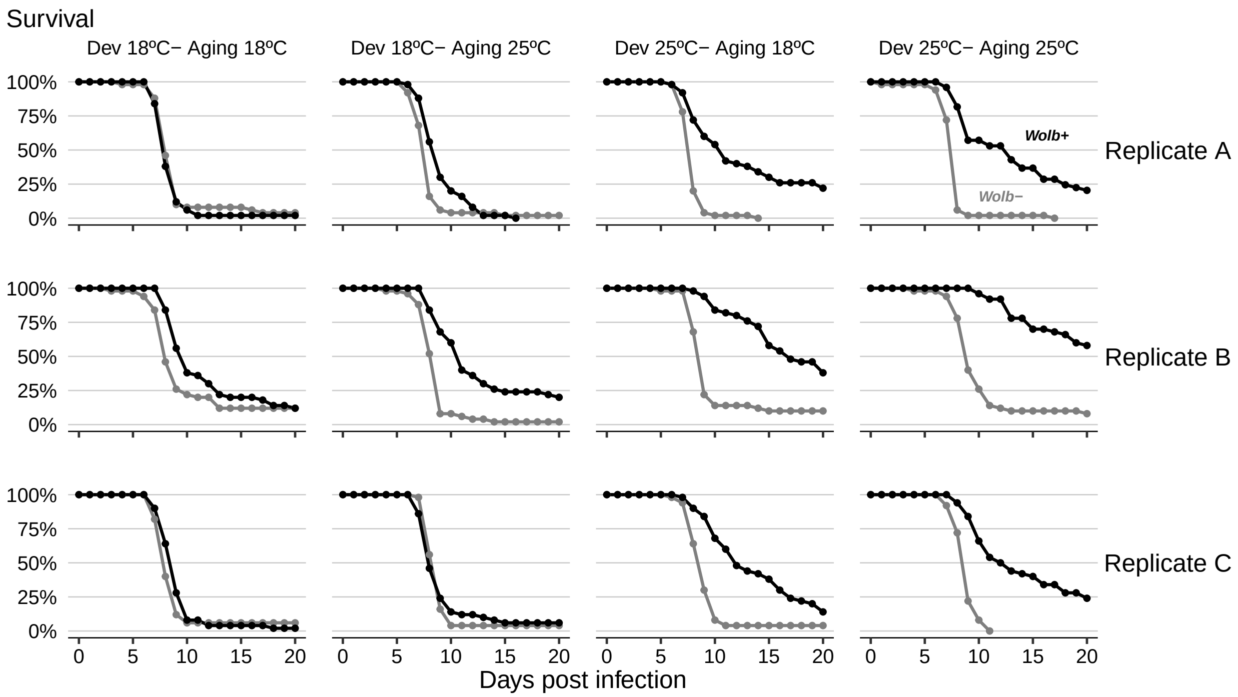

### Figure 4 and S5

#### DCV levels at early infection

```
DCV_early <- fread("dataset_s10.txt")[,lapply(.SD,char_asfactor)]
DCV_early <- filter(DCV_early, Treatment == "DCV")
DCV_early$logDCV <- ifelse(is.na(DCV_early$rel_DCV),log10(min(DCV_early$rel_DCV,na.rm = T)/10),log10(DCV
DCV_early$Time <- as.factor(as.character(DCV_early$Time))
```

```
#linear model
lm_DCV_early <- lm(logDCV~Wolb*Time, data=DCV_early)
Anova(lm_DCV_early)
```

```
## Anova Table (Type II tests)
##
## Response: logDCV
##             Sum Sq Df  F value    Pr(>F)
## Wolb        26.662  1  63.4876 2.462e-10 ***
## Time       288.807  5 137.5440 < 2.2e-16 ***
## Wolb:Time   15.928  5   7.5856 2.571e-05 ***
## Residuals   20.158 48
## ---
## Signif. codes:  0 '***' 0.001 '**' 0.01 '*' 0.05 '.' 0.1 ' ' 1
```

```
summary(lm_DCV_early)
```

```
##
## Call:
## lm(formula = logDCV ~ Wolb * Time, data = DCV_early)
```

```
##
## Residuals:
##      Min       1Q   Median       3Q      Max
## -2.06961 -0.29621 -0.01134  0.20753  1.55543
##
## Coefficients:
##                     Estimate Std. Error t value Pr(>|t|)
## (Intercept)         -1.66618    0.28981  -5.749 6.04e-07 ***
## WolbWolb+            0.29534    0.40985   0.721 0.474652
## Time0.25            -0.09621    0.40985  -0.235 0.815412
## Time0.5              2.17278    0.40985   5.301 2.87e-06 ***
## Time1                3.19419    0.40985   7.793 4.52e-10 ***
## Time2                5.49313    0.40985  13.403  < 2e-16 ***
## Time3                6.99756    0.40985  17.073  < 2e-16 ***
## WolbWolb+:Time0.25 -0.49534    0.57962  -0.855 0.397022
## WolbWolb+:Time0.5  -2.37547    0.57962  -4.098 0.000159 ***
## WolbWolb+:Time1    -1.79437    0.57962  -3.096 0.003273 **
## WolbWolb+:Time2    -2.28847    0.57962  -3.948 0.000257 ***
## WolbWolb+:Time3    -2.81762    0.57962  -4.861 1.29e-05 ***
## ---
## Signif. codes:  0 '***' 0.001 '**' 0.01 '*' 0.05 '.' 0.1 ' ' 1
##
## Residual standard error: 0.648 on 48 degrees of freedom
## Multiple R-squared:  0.9427, Adjusted R-squared:  0.9295
## F-statistic: 71.74 on 11 and 48 DF,  p-value: < 2.2e-16
```

There is an interaction between *Wolbachia* and time of infection

```
#compare effect of Wolb at the several time points
lsm_lm_DCV_early <- lsmeans::lsmeans(lm_DCV_early, pairwise~Wolb|Time,adj="none")
summary(lsm_lm_DCV_early,by=NULL,adj="holm")
```

```
## $lsmeans
##  Wolb  Time lsmean   SE df lower.CL upper.CL
##  Wolb- 0    -1.666 0.29 48   -2.538   -0.794
##  Wolb+ 0    -1.371 0.29 48   -2.243   -0.499
##  Wolb- 0.25 -1.762 0.29 48   -2.634   -0.890
##  Wolb+ 0.25 -1.962 0.29 48   -2.834   -1.090
##  Wolb- 0.5   0.507 0.29 48   -0.365    1.379
##  Wolb+ 0.5  -1.574 0.29 48   -2.446   -0.701
##  Wolb- 1     1.528 0.29 48    0.656    2.400
##  Wolb+ 1     0.029 0.29 48   -0.843    0.901
##  Wolb- 2     3.827 0.29 48    2.955    4.699
##  Wolb+ 2     1.834 0.29 48    0.962    2.706
##  Wolb- 3     5.331 0.29 48    4.459    6.203
##  Wolb+ 3     2.809 0.29 48    1.937    3.681
##
## Confidence level used: 0.95
## Conf-level adjustment: bonferroni method for 12 estimates
##
## $contrasts
##  contrast      Time estimate   SE df t.ratio p.value
##  Wolb- - Wolb+ 0      -0.295 0.41 48 -0.721  0.9493
```

```
##   Wolb- - Wolb+ 0.25    0.200 0.41 48  0.488  0.9493
##   Wolb- - Wolb+ 0.5     2.080 0.41 48  5.075  <.0001
##   Wolb- - Wolb+ 1       1.499 0.41 48  3.657  0.0019
##   Wolb- - Wolb+ 2       1.993 0.41 48  4.863  0.0001
##   Wolb- - Wolb+ 3       2.522 0.41 48  6.154  <.0001
##
## P value adjustment: holm method for 6 tests
```

```
log_DCV_early <- summary(lsm_lm_DCV_early,by=NULL,adj="holm")$contrasts[,3]
'^'(10,log_DCV_early)
```

```
## [1]   0.5065949   1.5848932 120.2627078  31.5522757  98.4306741 332.8744891
```

*Wolbachia* has a significant effect from 12h on.

##### Figure 4A

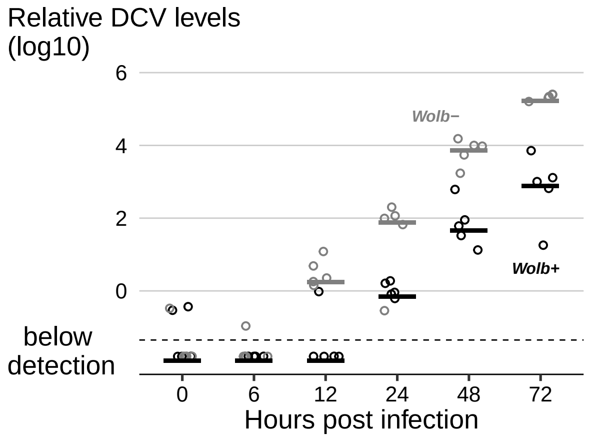

```
#another data set of early time points of DCV infection

DCV_early_doses <- fread("dataset_s11.txt")[,lapply(.SD,char_asfactor)]
DCV_early_doses$Time <- as.factor(as.character(DCV_early_doses$Time))
DCV_early_doses$logDCV <- ifelse(is.na(DCV_early_doses$Ratio),log10(min(DCV_early_doses$Ratio,na.rm = T)
```

```
#lm
lm_DCV_early_doses <- lm(logDCV~Genotype*Time*Dose, data=DCV_early_doses)
Anova(lm_DCV_early_doses)
```

```
## Anova Table (Type II tests)
##
## Response: logDCV
##                     Sum Sq Df F value    Pr(>F)
## Genotype            18.178  1 23.2119 2.473e-05 ***
## Time                55.014  1 70.2489 4.473e-10 ***
## Dose                16.738  1 21.3727 4.491e-05 ***
## Genotype:Time        0.790  1  1.0092  0.321608
## Genotype:Dose        0.775  1  0.9895  0.326331
## Time:Dose            7.165  1  9.1498  0.004505 **
## Genotype:Time:Dose   0.037  1  0.0469  0.829772
```

```
## Residuals          28.976 37
## ---
## Signif. codes:  0 '***' 0.001 '**' 0.01 '*' 0.05 '.' 0.1 ' ' 1
```

```
lm_DCV_early_doses_simple <- lm(logDCV~Genotype+Time*Dose, data=DCV_early_doses)
Anova(lm_DCV_early_doses_simple)
```

```
## Anova Table (Type II tests)
##
## Response: logDCV
##           Sum Sq Df F value    Pr(>F)
## Genotype  18.178  1 23.7496 1.771e-05 ***
## Time      54.720  1 71.4915 1.929e-10 ***
## Dose      16.611  1 21.7026 3.490e-05 ***
## Time:Dose  6.981  1  9.1213  0.004387 **
## Residuals 30.616 40
## ---
## Signif. codes:  0 '***' 0.001 '**' 0.01 '*' 0.05 '.' 0.1 ' ' 1
```

```
summary(lm_DCV_early_doses_simple)
```

```
##
## Call:
## lm(formula = logDCV ~ Genotype + Time * Dose, data = DCV_early_doses)
##
## Residuals:
##     Min      1Q  Median      3Q     Max
## -2.4281 -0.2828  0.1308  0.4261  1.3118
##
## Coefficients:
##                 Estimate Std. Error t value Pr(>|t|)
## (Intercept)       0.7896     0.2844   2.776 0.008330 **
## GenotypewMelCS  -1.2750     0.2616  -4.873 1.77e-05 ***
## Time24           1.4349     0.3654   3.927 0.000331 ***
## DoseE9           0.4480     0.3654   1.226 0.227349
## Time24:DoseE9    1.5771     0.5222   3.020 0.004387 **
## ---
## Signif. codes:  0 '***' 0.001 '**' 0.01 '*' 0.05 '.' 0.1 ' ' 1
##
## Residual standard error: 0.8749 on 40 degrees of freedom
## Multiple R-squared:  0.7593, Adjusted R-squared:  0.7352
## F-statistic: 31.54 on 4 and 40 DF,  p-value: 6.901e-12
```

*Wolbachia* confers significant resistance at these early time points

**Figure S5A**

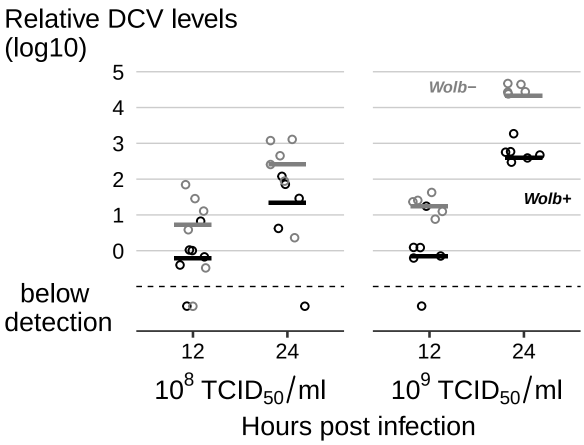

#### FHV levels at early infection

```
FHV_early <- fread("dataset_s12.txt")[,lapply(.SD,char_asfactor)]
FHV_early <- filter(FHV_early, Treatment == "FHV")
FHV_early$logFHV<- ifelse(is.na(FHV_early$rel_FHV),log10(min(FHV_early$rel_FHV,na.rm = T)/10),log10(FHV_
FHV_early$Time <- as.factor(as.character(FHV_early$Time))
```

```
#linear model
lm_FHV_early <- lm(logFHV~Wolb*Time, data=FHV_early)
Anova(lm_FHV_early)
```

```
## Anova Table (Type II tests)
##
## Response: logFHV
##            Sum Sq Df  F value    Pr(>F)
## Wolb        1.123  1  42.3198 2.428e-06 ***
## Time       95.652  4 901.1750 < 2.2e-16 ***
## Wolb:Time   0.421  4   3.9703   0.01571 *
## Residuals   0.531 20
## ---
## Signif. codes:  0 '***' 0.001 '**' 0.01 '*' 0.05 '.' 0.1 ' ' 1
```

```
summary(lm_FHV_early)
```

```
##
## Call:
## lm(formula = logFHV ~ Wolb * Time, data = FHV_early)
##
## Residuals:
##      Min       1Q   Median       3Q      Max
## -0.27644 -0.05261  0.00000  0.02928  0.41231
##
## Coefficients:
##                     Estimate Std. Error t value Pr(>|t|)
## (Intercept)       -1.247e+00  9.405e-02 -13.257 2.29e-11 ***
## WolbWolb+          3.430e-16  1.330e-01   0.000 1.000000
## Time1              1.673e+00  1.330e-01  12.578 5.90e-11 ***
```

```
## Time2            2.757e+00  1.330e-01  20.731 5.44e-15 ***
## Time3            3.912e+00  1.330e-01  29.412  < 2e-16 ***
## Time6            5.352e+00  1.330e-01  40.240  < 2e-16 ***
## WolbWolb+:Time1 -4.261e-01  1.881e-01  -2.265 0.034751 *
## WolbWolb+:Time2 -7.455e-01  1.881e-01  -3.963 0.000766 ***
## WolbWolb+:Time3 -4.046e-01  1.881e-01  -2.151 0.043894 *
## WolbWolb+:Time6 -3.586e-01  1.881e-01  -1.906 0.071085 .
## ---
## Signif. codes:  0 '***' 0.001 '**' 0.01 '*' 0.05 '.' 0.1 ' ' 1
##
## Residual standard error: 0.1629 on 20 degrees of freedom
## Multiple R-squared:  0.9946, Adjusted R-squared:  0.9921
## F-statistic:   407 on 9 and 20 DF,  p-value: < 2.2e-16
```

There is a significant interaction between *Wolbachia* and time

```
#compare effect of Wolb at the several timepoints
lsm_lm_FHV_early <- lsmeans::lsmeans(lm_FHV_early, pairwise~Wolb|Time,adj="none")
summary(lsm_lm_FHV_early,by=NULL,adj="holm")
```

```
## $lsmeans
##  Wolb  Time lsmean    SE df lower.CL upper.CL
##  Wolb- 0    -1.247 0.094 20   -1.543   -0.950
##  Wolb+ 0    -1.247 0.094 20   -1.543   -0.950
##  Wolb- 1     0.426 0.094 20    0.130    0.723
##  Wolb+ 1     0.000 0.094 20   -0.297    0.297
##  Wolb- 2     1.510 0.094 20    1.214    1.807
##  Wolb+ 2     0.765 0.094 20    0.468    1.062
##  Wolb- 3     2.665 0.094 20    2.369    2.962
##  Wolb+ 3     2.261 0.094 20    1.964    2.557
##  Wolb- 6     4.105 0.094 20    3.809    4.402
##  Wolb+ 6     3.747 0.094 20    3.450    4.043
##
## Confidence level used: 0.95
## Conf-level adjustment: bonferroni method for 10 estimates
##
## $contrasts
##  contrast      Time estimate    SE df t.ratio p.value
##  Wolb- - Wolb+ 0       0.000 0.133 20 0.000   1.0000
##  Wolb- - Wolb+ 1       0.426 0.133 20 3.204   0.0178
##  Wolb- - Wolb+ 2       0.746 0.133 20 5.605   0.0001
##  Wolb- - Wolb+ 3       0.405 0.133 20 3.042   0.0193
##  Wolb- - Wolb+ 6       0.359 0.133 20 2.696   0.0278
##
## P value adjustment: holm method for 5 tests
```

```
log_FHV_early <- summary(lsm_lm_FHV_early,by=NULL,adj="holm")$contrasts[,3]
'^'(10,log_FHV_early)
```

```
## [1] 1.000000 2.667471 5.565594 2.538451 2.283328
```

*Wolbachia* has a small significant effect from 1 day on (2 to 6 fold)

**Figure S5B**

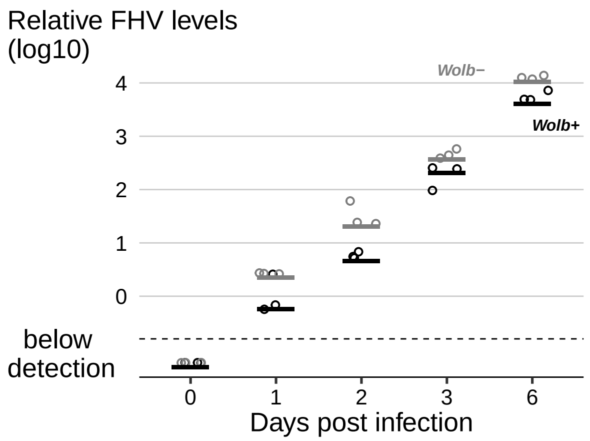

#### *Wolbachia* levels after antibiotics treatment

```
Antibiotics_levels <- fread("dataset_s13.txt")[,lapply(.SD,char_asfactor)]
Antibiotics_levels$logratio <- log10(Antibiotics_levels$ratio)
Antibiotics_levels$timepoint <- as.factor(as.character(Antibiotics_levels$timepoint))
Antibiotics_levels[,treatment:=relevel(treatment,"water")]
Antibiotics_treat <- filter(Antibiotics_levels, timepoint != "0")
```

```
#linear model
lmer_ant <- lmer(logratio~treatment*timepoint + (1|replicate), data=Antibiotics_treat)
Anova(lmer_ant)
```

```
## Analysis of Deviance Table (Type II Wald chisquare tests)
##
## Response: logratio
##                         Chisq Df Pr(>Chisq)
## treatment           1869.3483  5  < 2.2e-16 ***
## timepoint              0.1531  1     0.6956
## treatment:timepoint   29.3365  5  1.992e-05 ***
## ---
## Signif. codes:  0 '***' 0.001 '**' 0.01 '*' 0.05 '.' 0.1 ' ' 1
```

```
summary(lmer_ant)
```

```
## Linear mixed model fit by REML. t-tests use Satterthwaite's method [
## lmerModLmerTest]
## Formula: logratio ~ treatment * timepoint + (1 | replicate)
##    Data: Antibiotics_treat
##
## REML criterion at convergence: -116
##
## Scaled residuals:
##     Min      1Q  Median      3Q     Max
## -2.5493 -0.6249 -0.1255  0.5181  3.7513
##
```

```
## Random effects:
##  Groups    Name        Variance  Std.Dev.
##  replicate (Intercept) 0.0008164 0.02857
##  Residual              0.0207875 0.14418
## Number of obs: 155, groups:  replicate, 2
##
## Fixed effects:
##                                   Estimate Std. Error        df t value
## (Intercept)                        1.01457    0.04481  12.97522  22.642
## treatmentampicillin               -0.07155    0.05772 142.02299  -1.239
## treatmentethanol                   0.05314    0.05655 142.00000   0.940
## treatmentrifampicin               -0.93840    0.05655 142.00000 -16.594
## treatmentstreptomycin             -0.01454    0.05655 142.00000  -0.257
## treatmenttetracycline             -0.95028    0.05655 142.00000 -16.804
## timepoint30                        0.11133    0.05655 142.00000   1.969
## treatmentampicillin:timepoint30   -0.03037    0.08011 142.04098  -0.379
## treatmentethanol:timepoint30      -0.01597    0.07998 142.00000  -0.200
## treatmentrifampicin:timepoint30   -0.35046    0.07999 142.03508  -4.381
## treatmentstreptomycin:timepoint30 -0.11843    0.08081 142.01176  -1.466
## treatmenttetracycline:timepoint30 -0.20667    0.07998 142.00000  -2.584
##                                   Pr(>|t|)
## (Intercept)                       8.17e-12 ***
## treatmentampicillin                 0.2172
## treatmentethanol                    0.3490
## treatmentrifampicin                < 2e-16 ***
## treatmentstreptomycin               0.7975
## treatmenttetracycline              < 2e-16 ***
## timepoint30                         0.0509 .
## treatmentampicillin:timepoint30     0.7052
## treatmentethanol:timepoint30        0.8420
## treatmentrifampicin:timepoint30   2.28e-05 ***
## treatmentstreptomycin:timepoint30   0.1450
## treatmenttetracycline:timepoint30   0.0108 *
## ---
## Signif. codes:  0 '***' 0.001 '**' 0.01 '*' 0.05 '.' 0.1 ' ' 1
##
## Correlation of Fixed Effects:
##             (Intr) trtmntm trtmntth trtmntr trtmnts trtmnttt tmpn30 trtmntm:30
## trtmntmpcll -0.618
## tretmntthnl -0.631  0.490
## trtmntrfmpc -0.631  0.490   0.500
## trtmntstrpt -0.631  0.490   0.500    0.500
## trtmntttrcy -0.631  0.490   0.500    0.500   0.500
## timepoint30 -0.631  0.490   0.500    0.500   0.500   0.500
## trtmntmp:30  0.445 -0.721  -0.353   -0.353  -0.353  -0.353   -0.706
## trtmntth:30  0.446 -0.346  -0.707   -0.354  -0.354  -0.354   -0.707  0.499
## trtmntrf:30  0.446 -0.347  -0.353   -0.707  -0.353  -0.353   -0.707  0.499
## trtmntst:30  0.442 -0.343  -0.350   -0.350  -0.700  -0.350   -0.700  0.494
## trtmnttt:30  0.446 -0.346  -0.354   -0.354  -0.354  -0.707   -0.707  0.499
##             trtmntth:30 trtmntr:30 trtmnts:30
## trtmntmpcll
## tretmntthnl
## trtmntrfmpc
## trtmntstrpt
```

```
## trtmnttttrcy
## timepoint30
## trtmntmp:30
## trtmntth:30
## trtmntrf:30  0.500
## trtmntst:30  0.495       0.495
## trtmnttt:30  0.500       0.500      0.495
```

There is an interaction between treatment and timepoint

```r
#pairwise comparison of treatments at each time point
lsm_ant <- lsmeans::lsmeans(lmer_ant, pairwise~treatment|timepoint,adj="none")
summary(lsm_ant,by=NULL,adj="holm")
```

```
## $lsmeans
##  treatment    timepoint  lsmean     SE   df lower.CL upper.CL
##  water        10         1.0146 0.0448 13.0   0.8591   1.1700
##  ampicillin   10         0.9430 0.0463 14.6   0.7859   1.1001
##  ethanol      10         1.0677 0.0448 13.0   0.9122   1.2232
##  rifampicin   10         0.0762 0.0448 13.0  -0.0793   0.2316
##  streptomycin 10         1.0000 0.0448 13.0   0.8446   1.1555
##  tetracycline 10         0.0643 0.0448 13.0  -0.0912   0.2198
##  water        30         1.1259 0.0448 13.0   0.9704   1.2814
##  ampicillin   30         1.0240 0.0435 11.6   0.8694   1.1786
##  ethanol      30         1.1631 0.0448 13.0   1.0076   1.3185
##  rifampicin   30        -0.1630 0.0448 13.0  -0.3184  -0.0075
##  streptomycin 30         0.9929 0.0463 14.6   0.8358   1.1500
##  tetracycline 30        -0.0311 0.0448 13.0  -0.1865   0.1244
##
## Degrees-of-freedom method: kenward-roger
## Confidence level used: 0.95
## Conf-level adjustment: bonferroni method for 12 estimates
##
## $contrasts
##  contrast                    timepoint estimate     SE  df t.ratio p.value
##  water - ampicillin          10          0.0715 0.0577 142   1.239 1.0000
##  water - ethanol             10         -0.0531 0.0566 142  -0.940 1.0000
##  water - rifampicin          10          0.9384 0.0566 142  16.594 <.0001
##  water - streptomycin        10          0.0145 0.0566 142   0.257 1.0000
##  water - tetracycline        10          0.9503 0.0566 142  16.804 <.0001
##  ampicillin - ethanol        10         -0.1247 0.0577 142  -2.160 0.3248
##  ampicillin - rifampicin     10          0.8668 0.0577 142  15.014 <.0001
##  ampicillin - streptomycin   10         -0.0570 0.0577 142  -0.987 1.0000
##  ampicillin - tetracycline   10          0.8787 0.0577 142  15.220 <.0001
##  ethanol - rifampicin        10          0.9915 0.0566 142  17.533 <.0001
##  ethanol - streptomycin      10          0.0677 0.0566 142   1.197 1.0000
##  ethanol - tetracycline      10          1.0034 0.0566 142  17.743 <.0001
##  rifampicin - streptomycin   10         -0.9239 0.0566 142 -16.337 <.0001
##  rifampicin - tetracycline   10          0.0119 0.0566 142   0.210 1.0000
##  streptomycin - tetracycline 10          0.9357 0.0566 142  16.547 <.0001
##  water - ampicillin          30          0.1019 0.0555 142   1.835 0.6175
##  water - ethanol             30         -0.0372 0.0566 142  -0.657 1.0000
##  water - rifampicin          30          1.2889 0.0566 142  22.771 <.0001
```

```
##  water - streptomycin       30          0.1330 0.0577 142   2.303 0.2542
##  water - tetracycline       30          1.1569 0.0566 142  20.458 <.0001
##  ampicillin - ethanol       30         -0.1391 0.0555 142  -2.504 0.1744
##  ampicillin - rifampicin    30          1.1869 0.0555 142  21.369 <.0001
##  ampicillin - streptomycin  30          0.0311 0.0568 142   0.547 1.0000
##  ampicillin - tetracycline  30          1.0550 0.0555 142  18.994 <.0001
##  ethanol - rifampicin       30          1.3260 0.0566 142  23.427 <.0001
##  ethanol - streptomycin     30          0.1701 0.0577 142   2.947 0.0526
##  ethanol - tetracycline     30          1.1941 0.0566 142  21.115 <.0001
##  rifampicin - streptomycin  30         -1.1559 0.0578 142 -19.984 <.0001
##  rifampicin - tetracycline  30         -0.1319 0.0566 142  -2.331 0.2542
##  streptomycin - tetracycline 30         1.0240 0.0577 142  17.736 <.0001
##
## Degrees-of-freedom method: kenward-roger
## P value adjustment: holm method for 30 tests
```

*Wolbachia* levels in rifampicin and tetracycline flies significantly different from controls and other antibiotics at both time points ($p < 0.001$ for all). Rifampicin and tetracycline not significantly different at both time points ($p > 0.25$ for both). Other samples not significantly different from each other at each time point ($p > 0.05$ for all).

**Figures 4B and S5C**

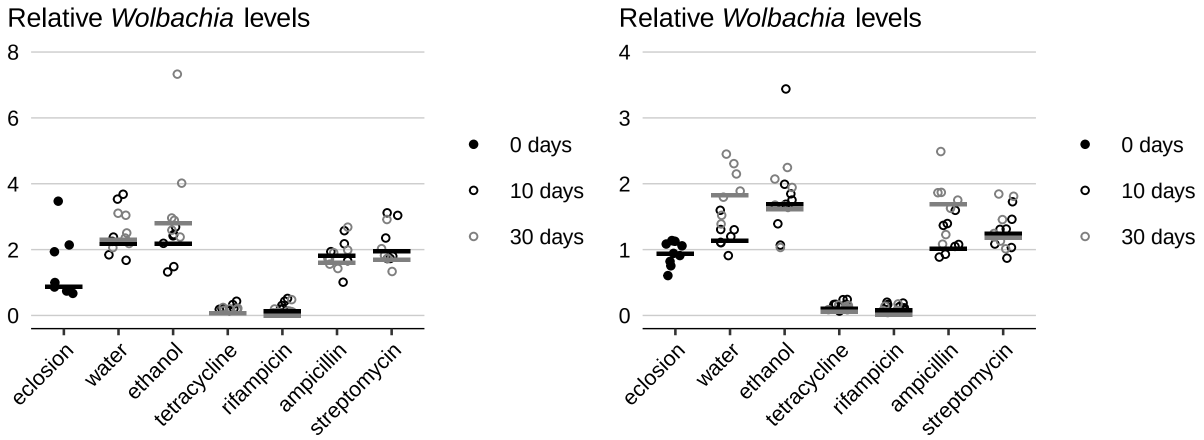

#### Survival to DCV after antibiotics

```
#load data
Survival_antibiotics <- fread("dataset_s14.txt")[,lapply(.SD,char_asfactor)]

Survival_antibiotics <- tidyr::separate(Survival_antibiotics, Condition, c("Wolbachia", "Treatment"), "
Survival_antibiotics <- tidyr::unite(Survival_antibiotics, RepFull, Treatment,Wolbachia,Test,Replicate,s

Survival_antibiotics <- Survival_antibiotics[,lapply(.SD,char_asfactor)]
Survival_antibiotics[,Treatment:=relevel(Treatment,"water")]
```

```r
#Diagnostics
#How many individuals were tested**
ftable(xtabs(~Test+Wolbachia+Treatment,Survival_antibiotics))
```

```
##                Treatment water ampicillin ethanol rifampicin streptomycin tetracycline
## Test Wolbachia
## A    CS                     50         50      50         50           50           50
##      iso                    50         50      50         50           50           49
## B    CS                     50         50      50         50           50           50
##      iso                    50         50      50         50           50           50
```

```r
##Data analysis
#Full model

Survival_antibiotics_cox<-coxme(Surv(Time,Status)~Wolbachia*Treatment+(1|RepFull)+(1|Test), Survival_an
Anova(Survival_antibiotics_cox,test.statistic = "LR")
```

```
## Analysis of Deviance Table (Type II tests)
##                     LR Chisq Df Pr(>Chisq)
## Wolbachia             70.955  1  < 2.2e-16 ***
## Treatment             41.817  5  6.416e-08 ***
## Wolbachia:Treatment   67.167  5  3.977e-13 ***
## ---
## Signif. codes:  0 '***' 0.001 '**' 0.01 '*' 0.05 '.' 0.1 ' ' 1
```

```r
summary(Survival_antibiotics_cox)
```

```
## Cox mixed-effects model fit by maximum likelihood
##   Data: Survival_antibiotics
##   events, n = 653, 1199
##   Iterations= 13 110
##                     NULL Integrated    Fitted
## Log-likelihood -4406.166  -4058.574 -3957.027
##
##                   Chisq   df p    AIC    BIC
## Integrated loglik 695.18 13.0 0 669.18 610.92
##  Penalized loglik 898.28 76.8 0 744.67 400.48
##
## Model:  Surv(Time, Status) ~ Wolbachia * Treatment + (1 | RepFull) +      (1 | Test)
## Fixed coefficients
##                                          coef   exp(coef)  se(coef)     z
## Wolbachiaiso                        2.6484521 14.13214648 0.4222592  6.27
## Treatmentampicillin                -0.5045895  0.60375339 0.5417512 -0.93
## Treatmentethanol                    0.4252938  1.53003993 0.4674664  0.91
## Treatmentrifampicin                 2.5392613 12.67030856 0.4202059  6.04
## Treatmentstreptomycin              -1.4907877  0.22519521 0.7079285 -2.11
## Treatmenttetracycline               2.5743570 13.12287695 0.4210857  6.11
## Wolbachiaiso:Treatmentampicillin    0.4011888  1.49359923 0.6407198  0.63
## Wolbachiaiso:Treatmentethanol      -0.2192497  0.80312117 0.5777168 -0.38
## Wolbachiaiso:Treatmentrifampicin   -2.4678268  0.08476888 0.5416011 -4.56
## Wolbachiaiso:Treatmentstreptomycin  1.3697108  3.93421287 0.7856511  1.74
## Wolbachiaiso:Treatmenttetracycline -2.5054517  0.08163871 0.5435895 -4.61
##                                          p
## Wolbachiaiso                       3.6e-10
## Treatmentampicillin                3.5e-01
```

```
## Treatmentethanol                      3.6e-01
## Treatmentrifampicin                   1.5e-09
## Treatmentstreptomycin                 3.5e-02
## Treatmenttetracycline                 9.7e-10
## Wolbachiaiso:Treatmentampicillin   5.3e-01
## Wolbachiaiso:Treatmentethanol      7.0e-01
## Wolbachiaiso:Treatmentrifampicin   5.2e-06
## Wolbachiaiso:Treatmentstreptomycin 8.1e-02
## Wolbachiaiso:Treatmenttetracycline 4.0e-06
##
## Random effects
##  Group   Variable  Std Dev    Variance
##  RepFull Intercept 0.65522621 0.42932138
##  Test    Intercept 0.29563700 0.08740124
```

There is an interaction between Treatment and *Wolbachia*

```
# Comparison of hazard ratios of Wolb versus no-Wolb at each treatment
mcp_Survival_antibiotics<-lsmeans::lsmeans(Survival_antibiotics_cox,pairwise~Wolbachia|Treatment)
summary(mcp_Survival_antibiotics,adj="holm",by=NULL)
```

```
## $lsmeans
##  Wolbachia Treatment    lsmean    SE  df asymp.LCL asymp.UCL
##  CS        water        -1.629 0.331 Inf    -2.576    -0.681
##  iso       water         1.020 0.239 Inf     0.336     1.703
##  CS        ampicillin   -2.133 0.387 Inf    -3.243    -1.024
##  iso       ampicillin    0.916 0.240 Inf     0.230     1.603
##  CS        ethanol      -1.203 0.297 Inf    -2.054    -0.353
##  iso       ethanol       1.226 0.236 Inf     0.550     1.901
##  CS        rifampicin    0.910 0.236 Inf     0.234     1.587
##  iso       rifampicin    1.091 0.239 Inf     0.408     1.774
##  CS        streptomycin -3.120 0.568 Inf    -4.748    -1.491
##  iso       streptomycin  0.899 0.238 Inf     0.216     1.581
##  CS        tetracycline  0.946 0.237 Inf     0.266     1.626
##  iso       tetracycline  1.089 0.242 Inf     0.396     1.781
##
## Results are given on the log (not the response) scale.
## Confidence level used: 0.95
## Conf-level adjustment: bonferroni method for 12 estimates
##
## $contrasts
##  contrast Treatment    estimate    SE  df z.ratio p.value
##  CS - iso water          -2.648 0.422 Inf -6.272  <.0001
##  CS - iso ampicillin     -3.050 0.477 Inf -6.388  <.0001
##  CS - iso ethanol        -2.429 0.388 Inf -6.261  <.0001
##  CS - iso rifampicin     -0.181 0.339 Inf -0.532  1.0000
##  CS - iso streptomycin   -4.018 0.659 Inf -6.099  <.0001
##  CS - iso tetracycline   -0.143 0.343 Inf -0.417  1.0000
##
## Results are given on the log (not the response) scale.
## P value adjustment: holm method for 6 tests
```

```r
#Wolb is protective in CTR, amp, and strp. Not in tet, rifa

contrast(mcp_Survival_antibiotics$contrasts,"pairwise",by="contrast")
```

```
## contrast = CS - iso:
##  contrast1                   estimate    SE  df z.ratio p.value
##  water - ampicillin            0.4012 0.641 Inf   0.626  0.9891
##  water - ethanol              -0.2193 0.578 Inf  -0.380  0.9990
##  water - rifampicin           -2.4678 0.542 Inf  -4.557  0.0001
##  water - streptomycin          1.3697 0.786 Inf   1.743  0.5029
##  water - tetracycline         -2.5055 0.544 Inf  -4.609  0.0001
##  ampicillin - ethanol         -0.6204 0.618 Inf  -1.003  0.9170
##  ampicillin - rifampicin      -2.8690 0.585 Inf  -4.903  <.0001
##  ampicillin - streptomycin     0.9685 0.817 Inf   1.186  0.8440
##  ampicillin - tetracycline    -2.9066 0.587 Inf  -4.952  <.0001
##  ethanol - rifampicin         -2.2486 0.516 Inf  -4.355  0.0002
##  ethanol - streptomycin        1.5890 0.768 Inf   2.070  0.3029
##  ethanol - tetracycline       -2.2862 0.518 Inf  -4.412  0.0001
##  rifampicin - streptomycin     3.8375 0.740 Inf   5.184  <.0001
##  rifampicin - tetracycline    -0.0376 0.484 Inf  -0.078  1.0000
##  streptomycin - tetracycline  -3.8752 0.742 Inf  -5.226  <.0001
##
## Results are given on the log (not the response) scale.
## P value adjustment: tukey method for comparing a family of 6 estimates
```

*Wolbachia* is protective in all conditions except in flies treated with tetracycline and rifampicin

##### Figures 4C and S5E

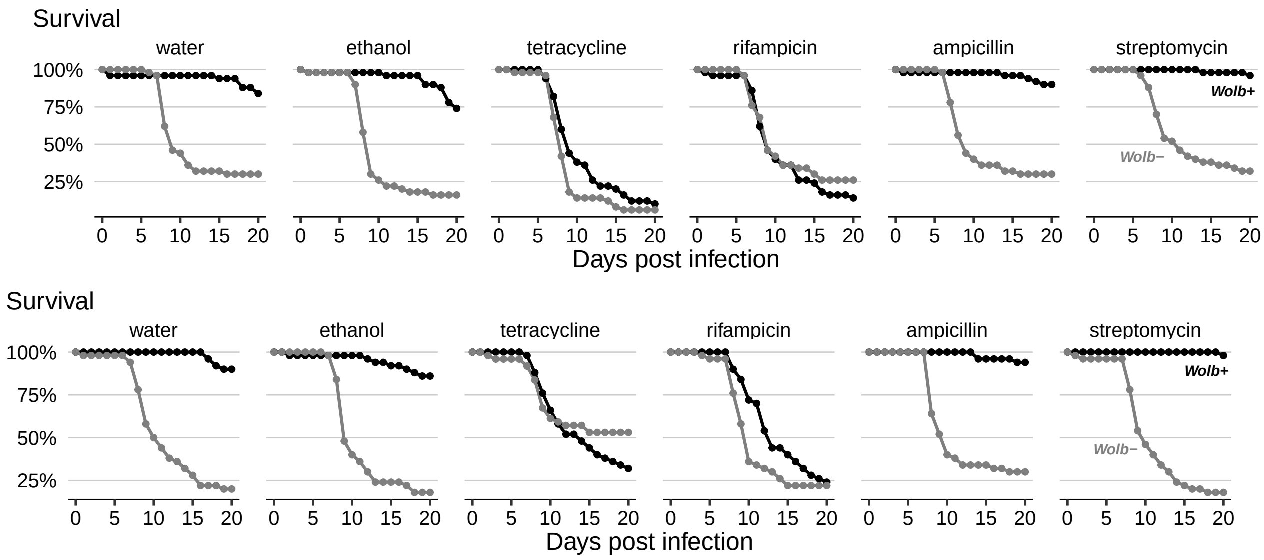

##### Bacteria levels in the gut after antibiotics

###### Figure S5D

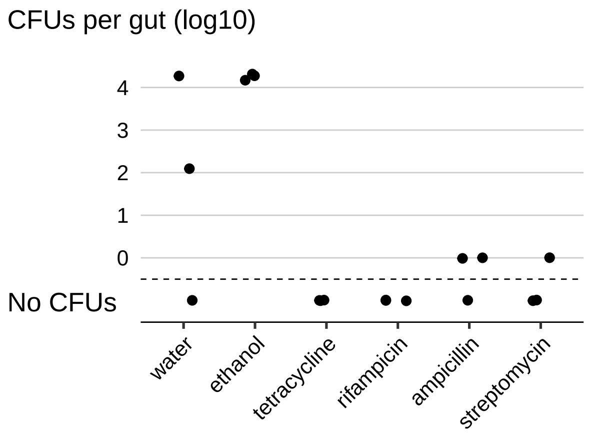

#### Session info

```
sessionInfo()
```

```
## R version 4.0.0 (2020-04-24)
## Platform: x86_64-apple-darwin17.0 (64-bit)
## Running under: macOS Catalina 10.15.5
##
## Matrix products: default
## BLAS:   /Library/Frameworks/R.framework/Versions/4.0/Resources/lib/libRblas.dylib
## LAPACK: /Library/Frameworks/R.framework/Versions/4.0/Resources/lib/libRlapack.dylib
##
## locale:
## [1] en_US.UTF-8/en_US.UTF-8/en_US.UTF-8/C/en_US.UTF-8/en_US.UTF-8
##
## attached base packages:
## [1] stats     graphics  grDevices utils     datasets  methods   base
##
## other attached packages:
##  [1] broom_0.5.6       lmerTest_3.1-2    lme4_1.1-23       Matrix_1.2-18
##  [5] reshape2_1.4.4    car_3.0-7         carData_3.0-3     multcomp_1.4-13
##  [9] TH.data_1.0-10    MASS_7.3-51.5     mvtnorm_1.1-0     lsmeans_2.30-0
## [13] emmeans_1.4.6     data.table_1.12.8 coxme_2.2-16      bdsmatrix_1.3-4
## [17] survival_3.1-12   lemon_0.4.4       forcats_0.5.0     stringr_1.4.0
## [21] dplyr_0.8.5       purrr_0.3.4       readr_1.3.1       tidyr_1.0.2
## [25] tibble_3.0.1      ggplot2_3.3.0     tidyverse_1.3.0   plyr_1.8.6
##
## loaded via a namespace (and not attached):
##  [1] nlme_3.1-147        pbkrtest_0.4-8.6    fs_1.4.1
##  [4] lubridate_1.7.8     httr_1.4.1          numDeriv_2016.8-1.1
##  [7] tools_4.0.0         backports_1.1.6     R6_2.4.1
## [10] DBI_1.1.0           colorspace_1.4-1    withr_2.2.0
## [13] tidyselect_1.0.0    gridExtra_2.3       curl_4.3
## [16] compiler_4.0.0      cli_2.0.2           rvest_0.3.5
```

```
## [19] xml2_1.3.1         sandwich_2.5-1     labeling_0.3
## [22] scales_1.1.0       digest_0.6.25      foreign_0.8-78
## [25] minqa_1.2.4        rmarkdown_2.1      rio_0.5.16
## [28] pkgconfig_2.0.3    htmltools_0.4.0    dbplyr_1.4.3
## [31] rlang_0.4.5        readxl_1.3.1       rstudioapi_0.11
## [34] farver_2.0.3       generics_0.0.2     zoo_1.8-7
## [37] jsonlite_1.6.1     zip_2.0.4          magrittr_1.5
## [40] Rcpp_1.0.4.6       munsell_0.5.0      fansi_0.4.1
## [43] abind_1.4-5        lifecycle_0.2.0    stringi_1.4.6
## [46] yaml_2.2.1         grid_4.0.0         parallel_4.0.0
## [49] crayon_1.3.4       lattice_0.20-41    haven_2.2.0
## [52] splines_4.0.0      hms_0.5.3          knitr_1.28
## [55] pillar_1.4.3       boot_1.3-24        estimability_1.3
## [58] codetools_0.2-16   reprex_0.3.0       glue_1.4.0
## [61] evaluate_0.14      modelr_0.1.7       vctrs_0.2.4
## [64] nloptr_1.2.2.1     cellranger_1.1.0   gtable_0.3.0
## [67] assertthat_0.2.1   xfun_0.13          openxlsx_4.1.4
## [70] xtable_1.8-4       statmod_1.4.34     ellipsis_0.3.0
```
